# Sparse deconvolution of cell type medleys in spatial transcriptomics

**DOI:** 10.1101/2025.02.07.637009

**Authors:** Nuray Sogunmez Erdogan, Deniz Eroglu

## Abstract

Mapping cell distributions across spatial locations with whole-genome coverage is essential for understanding cellular responses and signaling pathways. However, current deconvolution models often assume strong overlap between reference and spatial datasets, neglecting biological constraints like sparsity and cell-type variations. As a result, these methods rely on brute-force algorithms that ignore tissue complexity, leading to inaccurate predictions, particularly in heterogeneous or unmatched datasets.

We introduce Weight-Induced Sparse Regression (WISpR), a machine learning algorithm that integrates spot-specific hyperparameters and sparsity-driven modeling. Unlike brute-force methods, WISpR accurately predicts cell-type distributions while maintaining biological coherence, even in unmatched datasets. Benchmarking against five leading methods across ten datasets, WISpR consistently outperformed competitors and predicted cellular landscapes in both normal and cancerous tissues.

By leveraging sparse cell-type arrangements, WISpR provides biologically informed, high-resolution cellular maps. Its ability to decode tissue organization in both healthy and diseased states marks a major advancement in spatial transcriptomics, setting a new standard for accurate deconvolution.

## 1 Background

*Biological coherence*, the harmonious integration of cellular components and their interactions, underpins the complex organization and functionality of living systems. In nature, species form organizations that serve to perpetuate their lineage or enhance the efficiency of community tasks while minimizing energy consumption at different levels of life. Similarly, at the genetic level, coherence in region-specific gene expression allows cells to perform specialized roles, contributing to tissue complexity and organismal physiology[1–6]. Despite this, variability within cell types introduces *heterogeneity*, a key feature of complex tissues. Our understanding of these complex biological systems depends on our ability to accurately infer deconvolutions from observations. Yet, this coherence and heterogeneity dilemma becomes even more noticeable when the datasets originate from different sources. In such cases, the reference profiles may not adequately represent the cell types present in the spatial data, as the profiles of similar cell types can vary due to biological or temporal changes. However, the global goal is to establish a comprehensive collection of cellular transcriptomics profiles that can be used to map all possible cells in tissues and accurately detect disease states and locations. Therefore, accurately deconvoluting cell types from large libraries derived from various studies and understanding their interactions and organizations in spatial contexts is crucial for unraveling cellular heterogeneity and tissue complexity. Consequently, understanding the cellular landscapes and interactions opens up possibilities for deciphering complex biological processes such as cellular development, metabolism, and signal transduction[7–9] and diseases such as cancer, neurological disorders, and others[10–12].

*Spatial transcriptomics*, deciphering the location of gene expression profiles, allows us to identify tissue regions with transcriptional differences between capture spots. However, capture spots may contain multiple cell types or fractions of cells (Visium, Visium-HD, Tomo-Seq, Slide-Seq)[13–15] or subcellular spatial expression data with high dropout rates, producing sparse count matrices (as seen in Stereo-Seq, Seq-Scope, PIXEL-Seq)[16–18]. Consequently, directly extracting the whole genome coverage at single-cell resolution is impossible with current spatial transcriptomics technologies. On the other hand, *single-cell RNA sequencing* (scRNA-Seq) technology has revolutionized the study of cellular heterogeneity and gene expression at single-cell resolution, allowing examination of cell-specific transcriptomes within tissues. Despite potential confounders and biological variations[19–21], scRNA-Seq has revealed previously inaccessible insights into cellular diversity and function[22–25]. However, scRNA-Seq lacks spatial information on cell types’ original tissue locations. Hence, a combinatorial deconvolution approach using the cell-type information from scRNA-Seq and the spatial information from spatial transcriptomics is a natural solution to the problem.

Accurate predictions of cell types from spatial capture spots require deconvolution methods that account for biological constraints, including sparsity and non-negative presence of cells. Therefore, the availability of scRNA-Seq data and its direct utilization to deconvolute cell types in spatial transcriptomics are insufficient to accurately determine the correct cell distributions within the target tissue section. Existing deconvolution tools aim to integrate scRNA-Seq and spatial transcriptomics to create tissue atlases[26–31]. However, many methods neglect biological constraints, forcing predictions of all cell types across spatial locations, often resulting in biologically implausible outcomes, obscuring spatial organization, and hindering the discovery of critical disease mechanisms rooted in cellular communication. Therefore, finding an optimal and biologically reliable deconvolution algorithm for accurate mapping of cell types in spatial transcriptomics data remains an open problem.

The challenge lies in developing a computational tool that effectively aligns spatial transcriptomics with scRNA-Seq data while accounting for biological constraints to discard undesired cells during the deconvolution process. Deconvolution is often approached using regression methods to solve the linear equation ***y*** = ***Ax***. However, the high coherence of gene expressions among different cell types can pose difficulties for traditional regression approaches, as the column vectors in matrix ***A*** may be very similar. Attempts are made to overcome the problem by relying on correlation-based thresholding to select cell-type marker genes, assuming that these gene sets are highly specific to certain cell types [32]. Yet, in practice, some marker genes can be expressed in multiple cell types, i.e. coherent, or show variable expression depending on the context, which can reduce the specificity and reliability of cell-type identification. Furthermore, differentiating between closely related cell states or subtypes that do not possess distinct marker genes, potentially leading to inaccuracies in estimating cell-type proportions. Given the nature of our problem, the solution is expected to identify only relevant cell types, which are a few in reality, per spot. Therefore, sparse recovery methods (also known as compressed sensing or basis pursuit) emerge as clear candidates. Sparse deconvolution aims to identify the most critical cell types in a mixture by eliminating the least significant ones, thereby facilitating the most accurate predictions.

This article presents the implementation of a machine learning-based algorithm called *Weight-Induced Sparse Regression (WISpR)*, which incorporates an intelligently configured sparsity property to eliminate irrelevant cells as required. WISpR is specifically designed to predict the optimal sparse arrangement of cell types within scRNA-Seq-derived spatial transcriptomic data and to generate cellular maps for various tissues.

WISpR addresses the limitations of existing methods by incorporating sparsity constraints and spot-specific hyperparameter and gene weight optimization, ensuring biologically coherent predictions even in noisy or unmatched datasets. In evaluations, WISpR demonstrated superior accuracy and sensitivity in four matched scenarios (where the reference scRNA-Seq and target spatial transcriptomics datasets originate from the same source) and six unmatched scenarios (where the reference and target datasets originate from different sources), using synthetic data derived from two distinct organisms and tissue types. Furthermore, WISpR provided biologically meaningful information by accurately mapping cell-type proportions and localizations in diverse tissues, including the developing human heart and a coronal mouse brain section. Its ability to resolve abundant and rare cell populations demonstrates its utility in studying tissue complexity and cellular organization. Moreover, WISpR accurately mapped cell states in breast cancer tissues, revealing regional cancer subtypes and clarifying the location of LumB traits in the DCIS regions. This precision highlights the potential of WISpR to uncover tumor heterogeneity, advancing our understanding of cancer progression and resistance mechanisms. By excelling in both synthetic benchmarks and real biological applications, WISpR sets a new standard for deconvolution, providing a versatile and reliable tool for studying tissue architecture and cellular dynamics in diverse contexts.

## 2 Results

### 2.1 Cellular heterogeneity and genetic coherence in reference scRNA-Seq datasets

Accurate prediction of cell types in spatial transcriptomics data (deconvolution) is essential to understand the spatial organization of cells that govern tissue functions. Today’s technology provides an ultimate technique called scRNA-Seq, which profiles cellular gene expressions and opens the possibility of applying deconvolution from scRNA-Seq. Deconvolution is commonly performed using regression methods that combine gene expressions for each cell type with specific multiplier constants[33]. Therefore, each cell type needs to be determined by clustering approaches[34–36], and represented by selected genes, such as differentially expressed genes (DEGs), which exhibit distinct expression levels or read counts across cell types or experimental conditions. Clustering approaches explore similarities and differences in multivariate data and draw inferences from unlabeled data, resulting in cell populations sharing similar characteristics compared to other data members. For example, 3,717 cells from scRNA-Seq data of a developing human heart were clustered into 15 cell types[37] and visualized in a 2D space with uniform model approximation and projection (UMAP)[38] (Fig. 1**a**). However, deconvolution of these cell types is non-trivial, since the standard regression methods are insufficient for the desired reconstruction due to the following two main facts:

- *Cellular heterogeneity within a single cell type*. Cell-to-cell variability within the same cell type reflects how biological systems are regulated, respond to external influences, and develop over time. The heterogeneous nature of cells in a single cluster is illustrated for Cluster-0 (Fig. 1**b**). The distance from the centroid (stars) to the maximum distance to the convex hull of the cluster (black line) illustrates the cellular dispersion *σ*_0_. Thus, the identification of DEGs, the expression values of which represent all cells found in Cluster-0, is not trivial to select accurate representative gene expressions (Fig. 1**b**).
- *Genetic coherence in differently annotated cell types*. Along with cellular heterogeneity in clusters, it is likely to observe coherent gene expressions in separately annotated cell types due to their biological or functional similarities (i.e., cellular subtypes of the same tissue). For instance, distinct cell types, such as Clusters-2, 3, and 4, are nested in the 2D UMAP projection due to cellular heterogeneity (Fig. 1**c**). In other words, while *σ_i_*, where *i* = 2, 3, and 4, represents the heterogeneity for Clusters-2, 3, and 4, the overlapping circles with radius *σ_i_*s illustrate the similarities. A pairwise Pearson correlation between the expression vectors ***x*** and ***y***, *ρ****_x_***,***_y_*** (see the Methods section for the Pearson correlation formula), quantitatively measures the similarities between DEGs (Fig. 1**d**). This result reveals how the transcript levels of the expression vectors for different cell types, such as cluster 3 and cluster 4 (*ρ*_3,4_ = 0.91), or cluster 1 and cluster 12 (*ρ*_1,12_ = 0.94), strongly correlate with each other, declaring possible reasons for the failure of deconvolution.

**Fig. 1.**
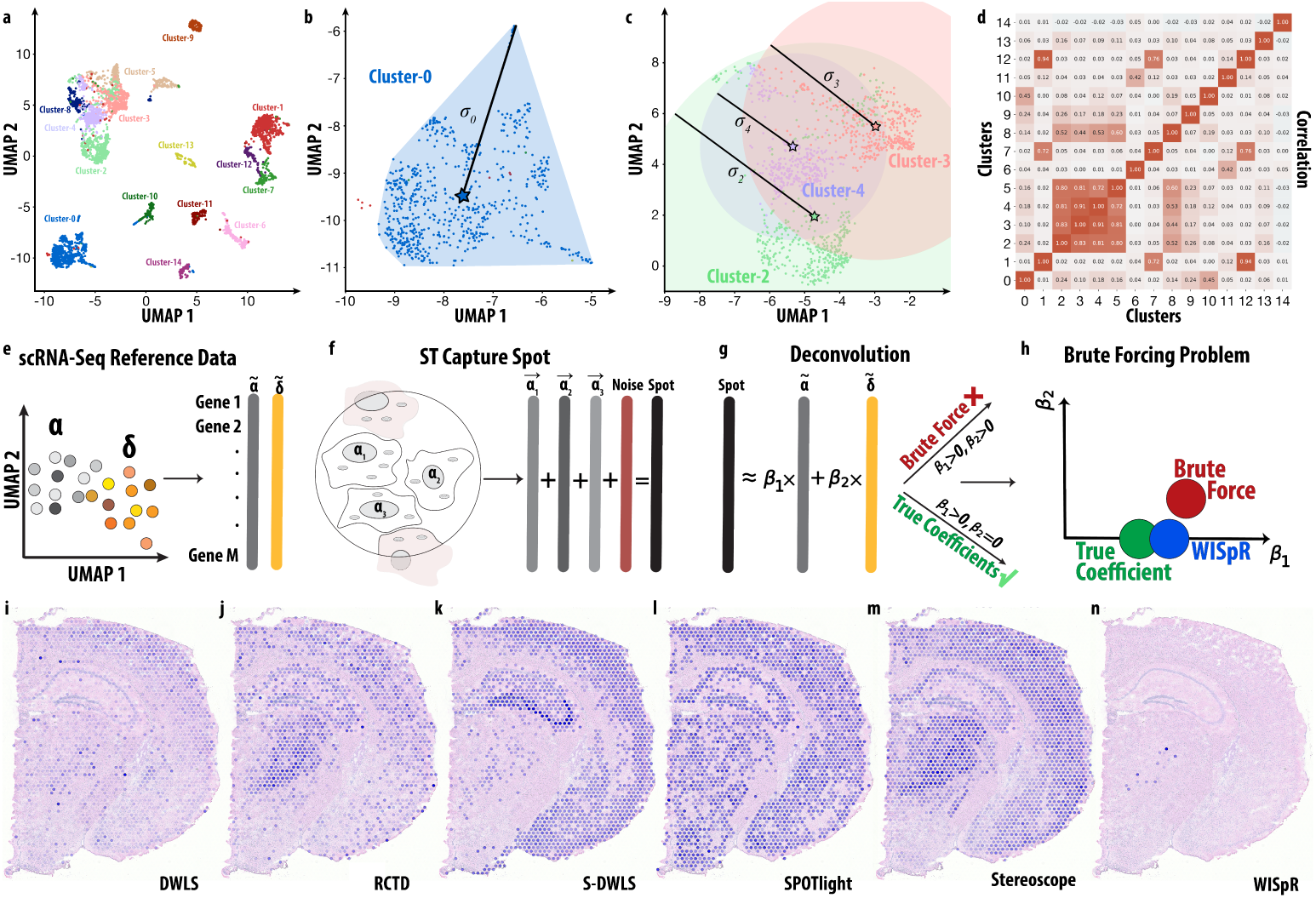
Deconvolution challenges due to coherence and heterogeneity. **a**, Two-dimensional UMAP representation of scRNA-Seq data from the developing human heart. Each color denotes cell types (clusters), revealing local and global structures based on their similarity. **b**, Illustration of high cellular heterogeneity within Cluster-0, represented by the line connecting the centroid (star) of Cluster-0 to the cell at the maximum distance. **c**, Overlapping clusters (Cluster-2 in green, Cluster-3 in orange, and Cluster-4 in blue) due to cell type similarity, with cluster radii calculated using centroid distances and cell locations at maximum distances. Large intersecting areas indicate similarity between cell types. **d**, Heatmap of Pearson’s correlations among clusters, with increased red intensity signifying higher correlations between functionally coherent cell types. The correlation values between clusters are displayed. **e**, The 2D UMAP plot of scRNA-Seq reference data showing cellular clusters from two cell types *α* (grey) and *δ* (orange) as separate clusters with their DEG vectors ***α̃*** and ***δ̃***. **f**, A spatial transcriptomics capture spot containing multiple cells from one cell type ***α*_1_**, ***α*_2_**, and ***α*_3_** (grey) and background noise (brick-red), illustrating the complexity of the spot’s transcriptomic profile. **g**, General deconvolution strategy using scRNA-Seq reference vectors to infer cell-type contributions in capture spots. Conventional brute-force methods assign non-negative coefficients, *β*_1_ and *β*_2_, to both cell types *α* and *δ* even when only *α* is present, leading to inaccurate, non-sparse solutions. **h**, Illustration of noise tolerance in deconvolution. Brute-force tools distribute coefficients across both cell types to account for noise, while WISpR uses thresholding to eliminate the contribution of *δ* and recover the true sparse coefficient for *α*, enhancing deconvolution accuracy. **i–n**, The reference scRNA-Seq data from the developing human embryonic heart is used to assess deconvolution approaches for distinguishing human heart-specific cell types from a mouse brain spatial transcriptomics dataset. Model predictions for atrial cardiomyocyte cell types in the mouse brain are presented. **i**, In DWLS prediction, human atrial cardiomyocytes show low percentages in mouse brain spots, with occurrences observed in cortical layers and thalamus. **j**, RCTD prediction reveals high percentages of cells in spots located in hippocampal, thalamic, and cortical regions. **k**, S-DWLS prediction, similar to RCTD, displays high percentages of cells in spots concentrated in cortical and hippocampal regions. **l**, SPOTlight and **m**, Stereoscope predicts a high abundance of cells across various brain zones, while **n**, WISpR predicts a minimal number of spots in the thalamus, supporting its more accurate prediction on mismatched datasets.

Therefore, both intra-cluster heterogeneity and inter-cluster coherence can yield computationally valid but biologically misleading regression fits.

Contemporary deconvolution techniques have been generated to overcome these challenges. These techniques can be categorized as model-based and model-free approaches depending on their proposed assumptions. The SPOTlight algorithm, a model-based approach, was among the first spatial deconvolution methods, using seeded non-negative matrix factorization followed by non-negative least squares regression to predict cell types in spatial positions[30]. Another widely used model-based algorithm, Robust Cell Types Decomposition (RCTD), on the other hand, relies upon to overcome the negative impacts of platform effects, Poisson sampling, and overdispersed counts on deconvolution[26]. The Stereoscope is a model-based negative binomial observation model[28] that is parameterized using logarithmic odds specific to the gene and cell types and a total number parameter specific to the gene. Estimated parameters will be used as input to the model to estimate the relative frequency of cells. However, the model-based algorithms are dependent on the selected model (which is accepted to be correct) and their training sets, which can cause unreliable results since the one-size-fits-all approach is, in general, inapplicable to all biological samples.

Among successful model-free approaches, the Dampened Weighted Least Squares (DWLS) method relies on overcoming the bias originating from the distribution of existing cell types or genes in transcriptomics data by employing bounded (dampened) weight multipliers to relatively scale the minimization error of the least squares regression[27]. Scaling is completed by an arbitrary function that reduces the error term to a relatively simpler one. However, this weight calculation, based on the inverse square of sequential solutions, may include negative values to be squared, which is biologically inaccurate. In addition, DWLS was originally designed to deconvolute bulk RNA sequencing datasets, making it computationally expensive to process thousands of capture spots found in spatial transcriptomics slides.

DWLS has then been further revised to the Spatial Dampened Weighted Least Squares (S-DWLS) algorithm, which attempts to improve prediction accuracy by incorporating cell type enrichment analysis[29]. S-DWLS is one of the first approaches to consider the existence of limited cell types in a single spot. However, similar to SPOTlight, RCTD, Stereoscope, and DWLS, S-DWLS adopts a generalized approach and neglects the effect of cellular heterogeneity and cell-type similarity, as explained above, by brute force fitting of the reference data to the spatial capture spots, overlooking the nuanced biological variability inherent in tissues.

Available, both model-based and model-free methods are designed to fit reference data to spatial capture spots, often without evaluating the true suitability of the reference, potentially leading to biased deconvolution. In other words, undesired cells’ or misestimated DEGs can be placed in the reference data. If a methodology uses a brute force approach to find the best fit, the result can contain unrelated cell types in the regression, which is generally overlooked in existing algorithms[26–30].

The challenges associated with spatial transcriptomics deconvolution when using the scRNA-Seq data as a reference is illustrated with a conceptual workflow in Fig. 1**e** - 1**h**. The 2D UMAP representation of the reference data (Fig. 1**e**) highlights the heterogeneous cluster of cell types annotated as *α* and *δ* as separate colors, and their DEG vectors ***α̃*** and ***δ̃*** with *M* number of genes, and *N* and *L* number of cells, where ***α̃*** = *f*(***α_i_***) for *i* = 1, 2*, .., N*, and ***δ̃*** = *f*(***δ_i_***) for *i* = 1, 2*, .., L*, respectively. The function *f* is an operation to find the representative vectors ***α̃*** and ***δ̃***, which is the mean value of each gene type within the cluster for our study. Furthermore, an example of a capture spot in spatial transcriptomics is given in Fig. 1**f**, where the spot is composed of three cells of a single cell type *α* along with contributions from background noise. The transcriptomic profile with *M* genes of the spot is thus a mixture of cells ***α_i_***, where *i* = 1, 2, 3 in this example, and noise, representing the measurement errors and fraction of cells partially contributing to the transcriptome profile of the spot. The general deconvolution strategy uses representative vectors, ***α̃*** and ***δ̃***, from the scRNA-Seq data to infer the cellular composition of the spatial capture spot. This approach relies on brute-force methods, assigning non-negative coefficients *β*_1_ and *β*_2_, to both cell types *α* and *δ*, respectively, even when only one cell type, *α*, is present in reality (Fig. 1**g**). This results in inaccurate predictions, as these methods fail to account for the sparsity of the true cellular composition. As a result, the concept of noise tolerance in deconvolution is required to predict the true coefficients. Brute-force tools attempt to explain the noise by indiscriminately assigning coefficients for almost all cell types given in reference data, which incorrectly represent different cell types, thereby introducing biologically implausible predictions that compromise interpretability. In contrast, WISpR is tailored to employ a thresholding mechanism to eliminate irrelevant cell types by assigning *β*_2_ = 0, and accurately assign sparse coefficients, *β*_1_ *>* 0, thus improving the precision of deconvolution in spatial data (Fig. 1**h**).

The real-life task is given to predict human heart cell types in a mouse brain tissue dataset. This setup, while representing an extreme case of dataset mismatch, highlights a fundamental criterion for algorithmic reliability: the ability to identify relevant cell types while discarding irrelevant ones. Here, probing atrial cardiomyocytes in a developing human heart (Cluster-7 in Fig. 1**a**) tested with alternative deconvolution methods (Fig. 1**e**–**i**, Fig. S1–S3), along with our model (Fig. 1**j**), to reveal incorrectly reconstructed mouse brain cells (10X Visium array) using human heart data[37], highlighting potential inaccuracies in disease studies where precise cell localization is critical. This outcome underscores a critical limitation: when tasked with even a straightforward dataset separation, these algorithms fail to exclude cell types absent from the target dataset.

While the given conceptual framework highlights the importance of accurate deconvolution, it initially assumes a highly non-ideal scenario where cell-type references have almost no match with the target tissue. This deliberate challenge was designed to test the robustness of alternative models under extreme conditions, using datasets from different organs and organisms. Although such mismatches are rare in biological practice, these tests are crucial for assessing model limitations. To complement this, we adopted a more biologically relevant approach, using adult mouse hippocampal scRNA-Seq data as a reference ([39]) (Fig.2**a**-2**c**) for deconvoluting hippocampus-only cells in postnatal day 8 (P8) [40] and adult mouse brain (10X Visium array). The objective is to ensure that reference hippocampal cells are located exclusively within the hippocampus, with no presence detected in other regions. This dual approach allowed us to rigorously evaluate the performance of the model in both extreme and realistic biological scenarios, consistently demonstrating the limitations of alternative models. An example is given here by using CA3 cells (Fig. 2**b**,2**c**). Similar to previous case, all tested alternative deconvolution methods (Fig. 2**d**–2**h** for P8 mouse, Fig. 2**j**–2**n** for adult mouse, and Figure. S4–S5), but WISpR (Fig. 2**i** for P8 mouse and Fig. 2**o** for adult mouse), over-reconstructed mouse brain cells using mouse hippocampus data. This outcome reveals that these algorithms are ineffective in removing absent cell types from the target data and tend to exaggerate the abundance of those present. Consequently, this limitation underscores the reliability of alternative tools in real-world applications where the ground truth may be unknown, and their tendency to over-represent present cell types while failing to exclude irrelevant ones could compromise accurate deconvolution. Therefore, methods that inherently sense biological phenomena, such as sparsity, intrinsically are needed to ensure deconvolution accuracy and to produce highly accurate high-resolution organ maps.

**Fig. 2.**
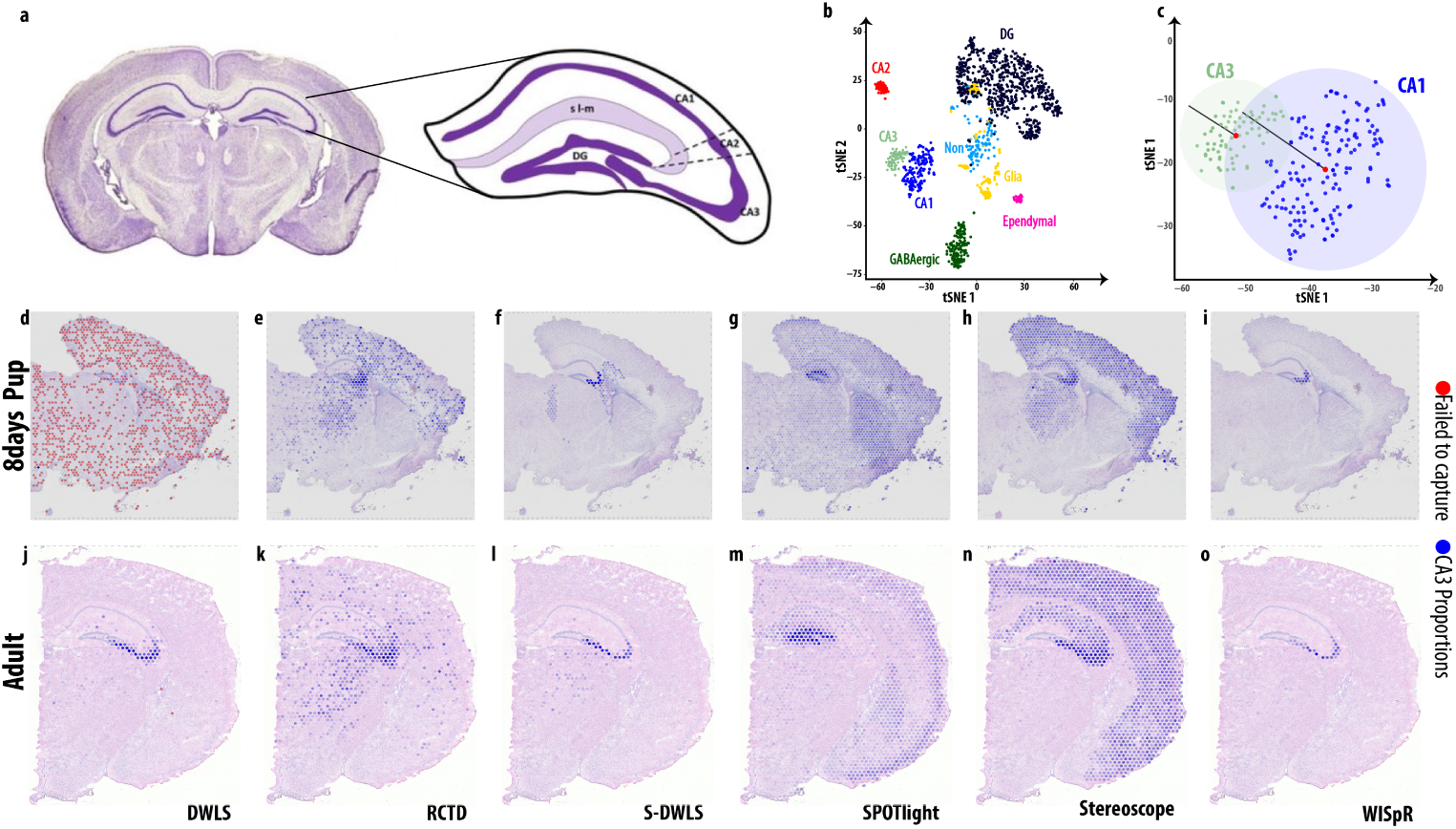
Overcoming limitations in sparse cell-type predictions for a biologically reliable task. **a**, Illustration of mouse hippocampus on mouse brain coronal section. **b**, 2D UMAP representation of the 8 HPF cell types. **c**, Overlapping CA1 and CA3 due to cell type similarity, with cluster radii calculated using centroid distances and cell locations at maximum distances. Large intersecting areas indicate similarity between cell types. **d-i**, Model predictions for CA3 in the P8 and **j-o** in adult mouse brain are presented. **d, j**, In DWLS prediction, CA3 cells are slightly over-represented in mouse brain spots (blue shade), with frequent spot prediction failures (red). **e, k**, RCTD prediction reveals high percentages of cells in spots located in hippocampal, thalamic and cortical regions. **f,l**, S-DWLS prediction displays confined but still disproportionate of cells in spots concentrated in stratum oriens, stratum pyramidale and stratum lucidum and thalamus regions. **g,m**, SPOTlight predicts a high abundance of CA3 cells in DG and across various brain zones including cortex. **h,n**, Stereoscope predicts CA3 in almost all hippocampus, thalamus and cortical regions, while **i,o** WISpR predicts CA3 cells in exactly CA3 morphological region in both P8 and adult mice, supporting its more accurate prediction on mismatched datasets.

### 2.2 WISpR: Weight-Induced Sparse Regression

Weight-Induced Sparse Regression (WISpR) introduces a game-changing approach to deconvolution, tailored for high-resolution datasets like scRNA-Seq. By integrating gene expression profiles with sparse regression, it overcomes the limitations of existing methods, including their inability to discard irrelevant cell types. Unlike traditional regression-based methods, WISpR dynamically adjusts for local sparsity by assigning spot-specific thresholds, a feature absent in comparable methods. This ensures that biologically irrelevant cell types are excluded from the deconvolution process. Therefore, it effectively addresses challenges such as local cell type sparsity, biological multicollinearity, cellular heterogeneity, gene expression imbalances, and potential overfitting during deconvolution.

*Sparsity* is a critical factor in cell type deconvolution, as tissue regions typically consist of only a limited number of cell types, each contributing to specific functional roles. This sparsity within larger cell populations necessitates a targeted approach, which WISpR delivers effectively. WISpR precisely promotes sparsity in transcriptomics data by systematically deactivating less impactful contributing cells during the deconvolution process, ensuring biologically plausible deconvolution. This innovation mitigates noise and enhances the reliability of cell-type identification, even in datasets with high variability or overlap. WISpR draws inspiration from the Sparse Identification of Nonlinear Dynamics approach, which achieves sparsity by sequentially reducing the function basis, setting their coefficients to zero if they fall below a specified threshold *τ* [41]. Consequently, WISpR excludes cell types from deconvolution if their contribution is deemed negligible.

WISpR excels at solving the challenging inverse deconvolution problem with a strong emphasis on achieving sparsity. This problem is is represented by ***y*** = ***Xβ***, where ***X*** = (***X***_1_*, . . .,* ***X****_Q_*) is an *M* × *Q* matrix and ***X****_i_* is an M-dimensional column vector, ***y*** = (***y***_1_*, . . .,* ***y****_L_*) is an *M* × *L*-dimensional matrix and ***y****_i_* is an *M* dimensional column vector, and ***β*** = (***β***_1_*, . . .* ***β****_Q_*) is a *Q*× *L*-dimensional matrix of coefficients and ***β****_i_* is a *Q*-dimensional column vector. In this equation, ***y****_i_* represents gene expressions in spatial transcriptomics spot-*i*, while ***X*** consists of gene expressions from various cell types obtained by scRNA-Seq data, organized as column vectors. The role of ***β****_i_* is to capture the coefficients that elucidate how the cell types within ***X*** contribute to the spatial transcriptomics data ***y****_i_*. The process of combining cell-type vectors in ***X*** with their corresponding coefficients in ***β*** results in the spatial transcriptomics matrix ***y***. Given that ***y*** is already known and does not require further manipulation, and the regression’s primary task is to determine the elements of ***β***, WISpR’s initial step focuses on creating a suitable matrix ***X***. This is achieved by leveraging differentially expressed genes (DEGs) from scRNA-Seq and spatial transcriptomics data, underscoring the importance of sparsity.

A DEG shows unique expression or read counts relative to other genes within specific cell types or conditions, serving as a cell type indicator. The presence of DEGs with observed expression levels distinguishes these cell types from others. However, capturing DEGs specific to cell types is a formidable task due to a range of challenges, including biological factors such as biological constraints and annotation, and methodological issues such as the absence of a gold standard scRNA-Seq dataset, high heterogeneity, dropout rates, potential errors in preprocessing, and the absence of biological ground truth, as depicted in Fig. 1**a**– 1**d**. Various methodological studies aim to accurately identify DEGs[42–45]. WISpR, on the other hand, identifies DEGs by considering the rarity of expression values of significant genes using an algorithm based on the Gini index[45, 46]. This algorithm assigns a score to each gene for each cell type (for detailed information on the DEG identification algorithm, please refer to the Methods section). The top 100 genes with the highest expression scores are selected as DEGs for each cell type from the scRNA-Seq data.

Cellular heterogeneity within a specific cell type can lead to an inadequate representation of DEGs from scRNA-Seq in spatial data. To address this, WISpR separates DEGs specific to cell types from scRNA-Seq data (Fig.3**a**,3**b**) and zone-specific DEGs from spatial data (Fig.3**c**,3**d**). Zone-specific DEGs characterize areas with similar transcriptional profiles. In contrast to S-DWLS, which assesses the enrichment of scRNA-Seq DEGs at each spot individually, WISpR combines scRNA-Seq DEGs with zone-specific DEGs (Fig. 3**e**). Consequently, WISpR infers DEGs and generates DEG vectors (***X****_i_*) for each cell type *i*. These vectors ***X****_i_* are assembled into a matrix, forming ***X*** for deconvolution (Fig. 3**f**) and are used to deconvolute the abundance and composition of cell types per spot (Fig. 3**g**).

**Fig. 3.**
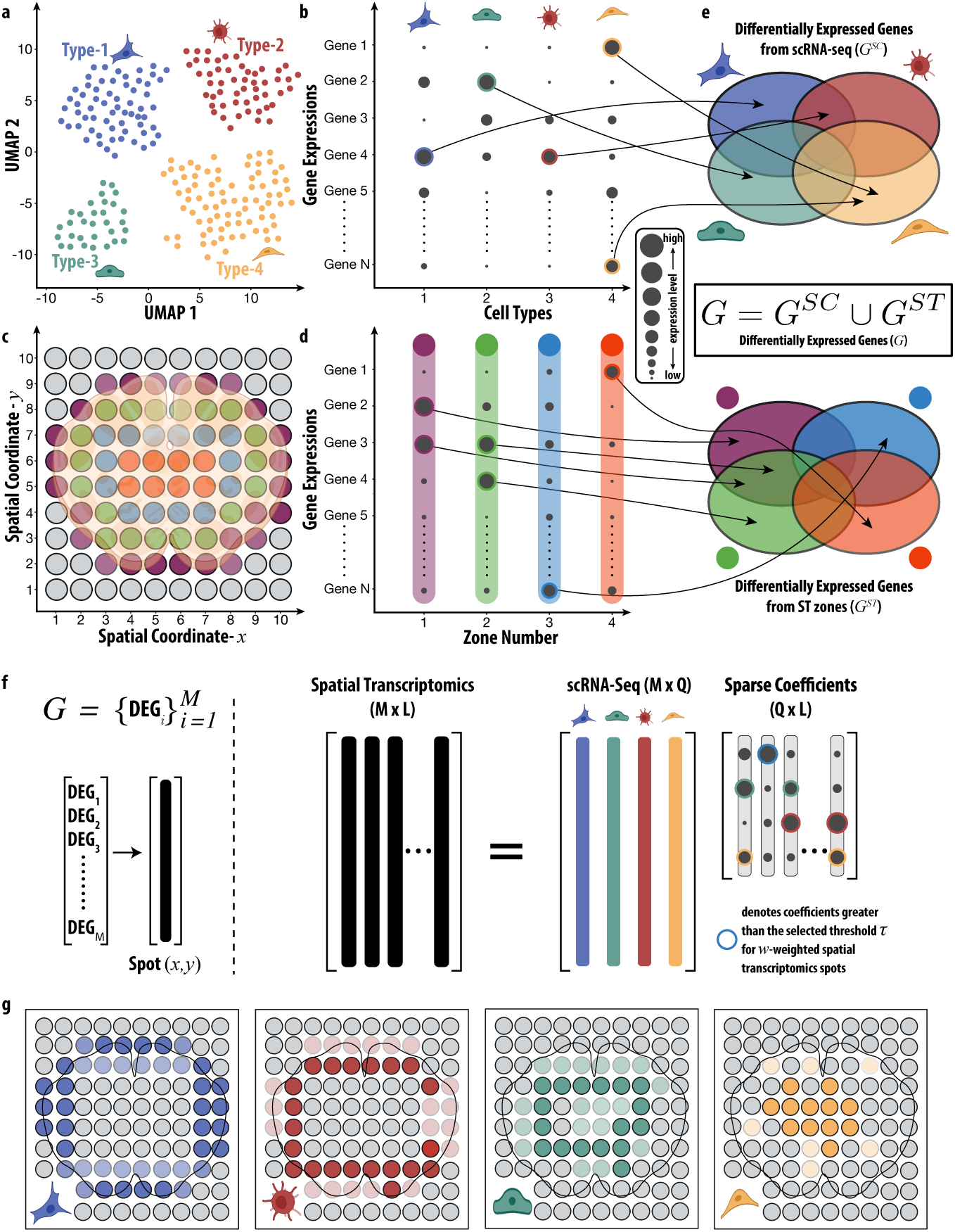
WISpR Spatial Deconvolution Workflow Overview. **a**, UMAP visualization of scRNA-Seq data showcasing four distinct cell types, each depicted in a unique color. **b**, Gene expression profiles for individual cell types derived from the intersection of genes (N) in scRNA-Seq and spatial transcriptomics datasets. **c**, Representative *x*- and *y*-coordinates of spots in the spatial transcriptomics dataset, with each color indicating a zone profile identified through clustering analysis. **d**, Gene expression profiles for individual spatial zones based on a selected gene set from the intersection of scRNA-Seq and spatial transcriptomics datasets. **e**, Identification of differentially expressed genes specific to cell types (*G^SC^*) and zones (*G^ST^*), defining cluster differentiation strength. The union of *G^SC^*and *G^ST^*forms the reference scRNA-Seq and spatial transcriptomics datasets (*G*) for WISpR deconvolution. **f**, WISpR algorithm summary. The spatial transcriptomics matrix (MxL) and reference scRNA-Seq matrix (MxQ) guide the prediction of the best sparse coefficient matrix (QxL), representing the number of genes (M) in *G* for each spot, number of spots (L), and number of reference cell types (Q). WISpR iteratively thresholds coefficients based on spot-specific weights (*w*) and thresholds (*τ*). **g**, Spatial deconvolution results reveal estimated abundance patterns, locations, and co-occurring cell type compositions for each reference cell type using the WISpR algorithm.

In practice, WISpR is a weight-induced sparse regression method that seeks the best and most sparse fit between scRNA-Seq and spatial transcriptomics. WISpR achieves this by iteratively eliminating negligible contributions of scRNA-Seq vectors ***X****_i_* in the basis matrix ***X***, setting *β_i,j_* = 0. The algorithm incorporates three crucial hyperparameters to account for biological heterogeneity in both the number of cell types and transcript levels in tissues: (*i*) spot-specific bias terms, (*ii*) spot-specific threshold parameters, and (*iii*) gene-specific weights per spot. Ridge regression, employing the Euclidean (*L*_2_) norm with penalization by adding a *bias term*, is used to reduce the standard error and improve the reliability of the regression. This discarding process iterates through *β_i,j_* values smaller than the specified *threshold*. Low-contributing vectors ***X****_i_* are removed by fixing *β_i,j_* = 0, and regression continues with an updated coefficient vector ***β****_i_* until no removable vectors remain in ***X***. However, unconditional removal of vectors can lead to undesirable outcomes, as the rarity of some cell types and expressed genes should be preserved to prevent false negative results. To address this, spot-specific *weight* parameters are introduced, derived from the initial prediction’s non-negative solution with the multiplicative inverse as the weight vector. Spot-specific bias, threshold, and weight parameters are optimized through an exhaustive search within a hyperparameter subset. This search algorithm simplifies WISpR’s usability, reducing the number of parameters while still identifying optimal parameters for each spot.

In summary, WISpR performs targeted deconvolution, retaining significant predictors while eliminating non-contributory ones. The enhanced cell-type sparsity achieved through this technology contributes to improved prediction accuracy in our model compared to alternative methods. A comprehensive description of mathematical methodology with a generalization to deconvolute all spatial spots in a matrix form (Fig. 3**f**) can be found in the Methods section.

### 2.3 Applications: Benchmarking

#### 2.3.1 Semi-Synthetic Data

Deconvolution in spatial transcriptomics typically assumes significant overlap between cell types in reference and spatial datasets. However, mismatched and unlabeled cell types often arise due to differences in data preparation, sequencing platforms, and protocols. Rare or closely related cell types in scRNA-Seq may remain indistinguishable from spatial transcriptomics data, adding technical variability. Additionally, spatial transcriptomics covers larger tissue sections, resulting in the inclusion of more cell types than are typically found in scRNA-Seq references. These challenges necessitate rigorous benchmarking of deconvolution models.

To evaluate WISpR’s robustness against these challenges, we tested it on semisynthetic spatial transcriptomics datasets generated using two distinct methodologies. Datasets were created following a Dirichlet distribution, simulating diverse cell ratios and capture location scenarios (as detailed in Andersson et al. (2020)[28] and the Methods section).

The first dataset was generated using human embryonic heart scRNA-Seq data[37], covering four scenarios: (1) dense sampling with 10 to 30 cells per capture spot, (2) sparse sampling (1 to 10 cells per spot, mimicking 10X Visium), (3) added noise, and (4) mislabeled data, each comprising 1,500 spots.

In the second approach, we combined cells from two sources—human embryonic heart[37] and mouse brain[47]—to test the impact of gene expression disparities and unmatched cell types on prediction performance. A blended dataset, denoted *Mix*[*a, b, c*], was generated to evaluate the deconvolution models. Here, *a* represents mouse brain cells common to reference scRNA-Seq and synthetic spatial datasets, *b* represents human heart cells, and *c* represents mouse brain cells absent in the spatial dataset. By design, *a* was fixed at 50%, while *b* + *c* = 50%. The expected outcome should exclude contributions from *b* and *c* if the deconvolution is reliable.

We compared WISpR to DWLS, RCTD, S-DWLS, Stereoscope, and SPOTlight using Root Mean Squared Error (RMSE) and F1 scores (detailed in the Methods section). Across the four scenarios, WISpR consistently achieved the lowest RMSE in Scenarios 1, 2, and 4, with values of 0.032 ± 0.023, 0.050 ± 0.049, and 0.057 ± 0.054, respectively. In Scenario 3, WISpR performed comparably to DWLS and RCTD (0.071 ± 0.075, 0.071 ± 0.072, and 0.068 ± 0.063, respectively) (Fig. 4**a**, 4**b**). Separate analysis of RMSE for *a*, *b*, and *c* in *Mix*[*a, b, c*] reveals the lowest errors in predicting true cells, along with minimal false positive rates for unmatched cells (Fig. S6). Statistical analysis using the Wilcoxon rank-sum test revealed that WISpR significantly outperformed competing models in Scenarios 1, 2, and 4 (*p <* 0.01) and did not show significant differences in RMSE distributions between DWLS (p = 0.258), RCTD (p = 0.740), and S-DWLS (p = 0.152) (Tables S1 and S2).

**Fig. 4.**
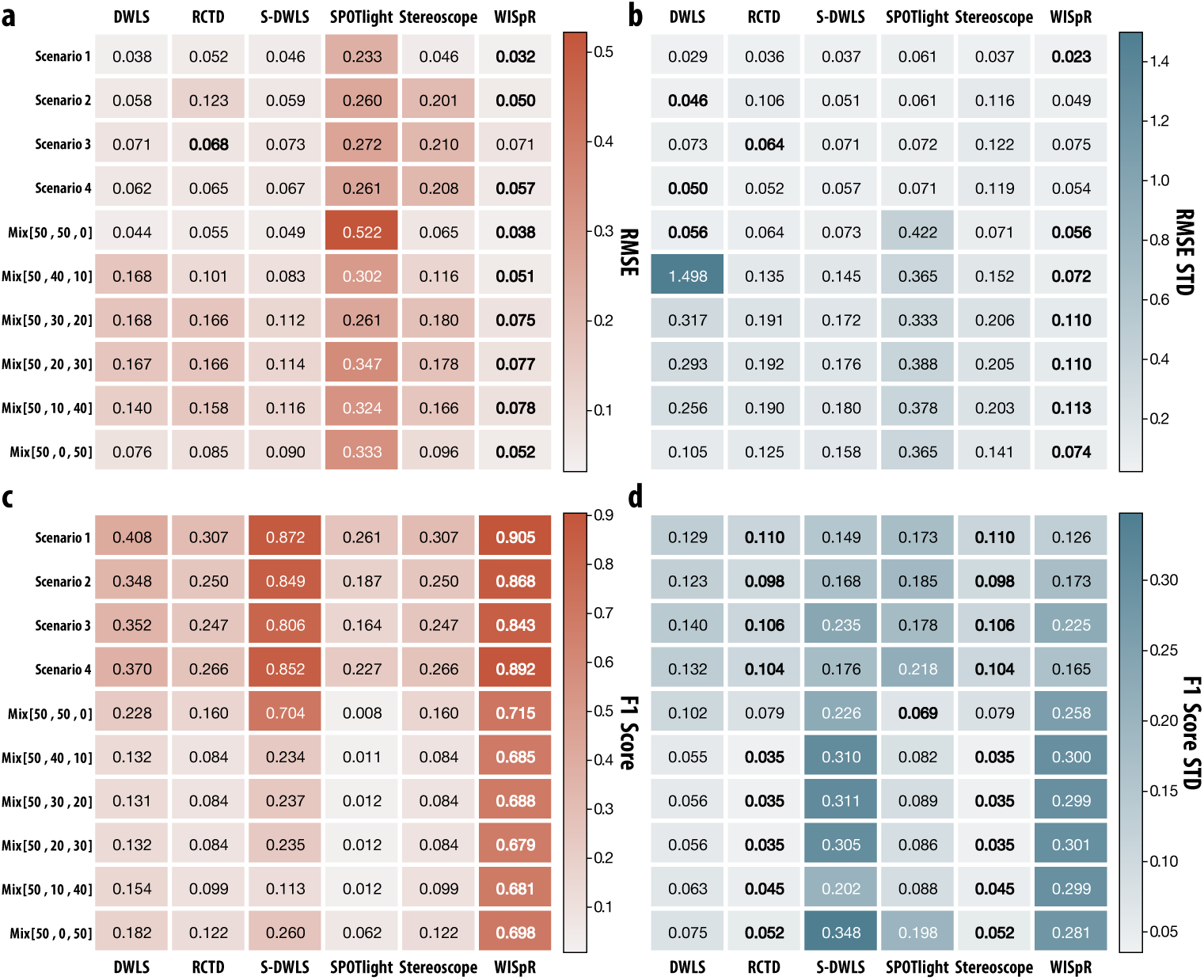
Overview of the comparative analysis of deconvolution model performances using synthetic spatial data in four different scenarios and six different mixture percentages. Four benchmark scenarios are generated by selected cells from 15 cell types in the developing human embryonic heart scRNA-Seq dataset. 6 different mixtures are generated using human heart cells and mouse brain cells. Tested deconvolution algorithms are DWLS, RCTD, S-DWLS, SPOTlight, Stereoscope, and WISpR. **a** A mean Root Mean Squared Error (RMSE) assessment of predictive performances of six methods’ performances. Except for Scenario 3, WISpR outperforms the other five models. In Scenario 3, a slightly better mean predictive performance was observed in RCTD model, where the distribution of errors between RCTD and WISpR displayed no statistical significance. **b**: The mean standard deviations of RMSE scores. WISpR emerged as low error prone especially in blended data (Mix[50,50,0], Mix[50,40,10], Mix[50,30,20], Mix[50,20,30], Mix[50,10,40], Mix[50,0,50]). **c** The mean F1 scores and **d** their mean standard deviations in all scenarios. When there are highly matching cell types between scRNA-Seq and synthetic data, WISpR’s and S-DWLS’s F1 scores are the highest; however, when unmatched cell types are introduced, the S-DWLS score decreases dramatically. The low standard deviations in DWLS, RCTD and SPOTlight indicate their ubiquitous false predictions of the non-existing cell types.

The RMSE distributions provide detailed insights into prediction errors beyond the mean RMSE, depicting the distribution of errors across six models in four scenarios. Computed with 20 bins that span the range between minimum and maximum error values, these distributions reveal distinctive patterns. Mean RMSE (*µ*), standard deviation (*σ*), and skewness (*γ*) values for each prediction are indicated on the plots (Fig. S19). WISpR, DWLS, RCTD, and S-DWLS exhibit left-skewed distributions, indicating high prediction accuracies, and Stereoscope displays left-skewed distributions solely for Scenario 1; meanwhile, a more bimodal distribution characterizes the increased errors for the remaining scenarios. In contrast, SPOTlight demonstrates right-skewed RMSE distributions in all four scenarios, indicative of higher predictive errors. Among the six models, WISpR consistently demonstrates satisfactory prediction accuracies based on its *µ*, *σ*, and *γ* scores, except Scenario 3, where the statistical calculations reveal no statistically significant differences between the RMSE distributions of WISpR and RCTD, and WISpR and DWLS. The red dashed lines denote the average RMSE values (*µ*), as indicated in the main Fig. 4.

F1 scores further highlighted the superior accuracy of WISpR. It consistently achieved scores exceeding 0.84 in all scenarios, demonstrating robust performance even under noisy and mislabeled conditions (Scenario 3, *F* 1 = 0.843 ± 0.225, Scenario 4, *F* 1 = 0.892 ± 0.165) (Fig. 4**c**, 4**d**). Individual F1 scores for *a* and total variational distances for *b* and *c* further exhibit robust prediction of WISpR independent of noise (Fig. S7). Notably, models such as RCTD and Stereoscope, while exhibiting low standard deviations, displayed reduced predictive precision due to consistently low F1 scores. In contrast, the higher F1 scores of WISpR underscore its ability to resolve subtle cellular patterns while minimizing false positives.

In blended datasets *Mix*[*a, b, c*], WISpR demonstrated outstanding predictive capacity, achieving the lowest RMSE values across all blends, with values ranging from 0.038 ± 0.056 for *Mix*[50, 50, 0] to 0.078 ± 0.113 for *Mix*[50, 0, 50], all of which are significantly lower (p<0.01) than the alternative model predictions (Fig. 4**a**, 4**b**, Table S2). These results highlight WISpR’s robustness across varying levels of mismatch and noise, where competing methods showed significant performance deterioration. Notably, S-DWLS, a widely used method, performed well in scenarios involving distinct cell types (i.e., *Mix*[50, 50, 0]), but struggled when similar cell types absent in the synthetic spatial dataset were introduced into the reference dataset. This difficulty underscores the challenges existing methods face when handling biologically similar cell types, particularly when some were shared between datasets and others were absent in the spatial data.

Moreover, the RMSE distributions illustrate the spread of prediction errors across six models in six blended datasets (Fig. S20). In this analysis, both the RMSE values for true positive cell-type predictions and the error rates for false positive cell-type predictions are computed. For DWLS, the error range falls within [0–30] for the *Mix*[50, 40, 10] blended data. To ensure a precise comparison of distributions and maintain similar distribution patterns within the range of [0,1], the distribution is plotted using 400 bins. Similarly, for the *Mix*[50, 30, 20] and *Mix*[50, 20, 30] datasets, the error rate ranges from 0 to 5.48, while for the *Mix*[50, 10, 40] dataset, it ranges from 0 to 4.69. Therefore, the distribution is plotted using 100 bins, with error values within the range of [0,1] depicted for comparison. For the remaining RMSE distributions, the number of bins is set to 20, covering the range between their minimum and maximum error values. The left skewness represents high accuracy in predictions. Across the six blended data, WISpR exhibited consistency in its mean RMSE (*µ*), standard deviation (*σ*), and skewness (*γ*) scores, where a high abundance of its predictions had the lowest error scores. The highest *γ* score of DWLS is attributed to its number of high prediction error values, therefore increasing the skewness of its distribution. RCTD, S-DWLS, and Stereoscope displayed left-skewed error distributions with a significantly lower number of good predictions compared to WISpR. SPOTlight resulted in bimodal error distributions, where a significant number of its predictions exhibited high errors. Furthermore, WISpR maintained high predictive accuracy in all tested configurations, with F1 scores consistently exceeding 0.68, even under challenging mismatch conditions. This performance reflects WISpR’s ability to accurately localize biologically relevant cell types while minimizing false-positive predictions. For example, in *Mix*[50, 50, 0], where 50% of the cells in the dataset were mouse brain cells and the remaining 50% were human heart cells, WISpR excelled at identifying the correct cell types with minimal errors (F1 = 0.715). Notably, S-DWLS also performed well in this configuration, achieving an F1 score of 0.704. However, its performance was dramatically reduced to 0.113 when similar mouse brain cells, absent in the synthetic spatial dataset, were intentionally introduced into other blended datasets. This highlights the lower sensitivity of S-DWLS to the incorporation of biologically similar cell types, which led to a decreased precision in distinguishing between cell types from different tissues, particularly in the presence of noise or additional reference mismatches (Fig. 4**c**,4**d**).

When comparing F1 values across all models (DWLS, RCTD, S-DWLS, SPOTlight, Stereoscope, and WISpR), WISpR consistently outperforms the alternatives, achieving substantial improvements in prediction accuracy. In all datasets, WISpR achieves F1 scores above 0.68, with the highest score of 0.715, while competing methods such as SPOTlight, Stereoscope and S-DWLS produce significantly lower scores in most cases, often below 0.26. For instance, compared to S-DWLS, which performs well in *Mix*[50, 50, 0], but struggles in other configurations, WISpR demonstrates an improvement of up to 502% (*Mix*[50, 10, 40]). Similarly, compared to DWLS, which shows a maximum F1 score of 0.228, WISpR offers an enhancement of approximately 213%. Even Stereoscope, the closest competitor in certain configurations, lags significantly behind, highlighting WISpR’s unparalleled robustness and predictive capacity (Fig. 4**c**, 4**d**).

WISpR’s performance in blended datasets demonstrates its capability to account for biological sparsity and natural coherence, even in cross-tissue or cross-species analyses. The method’s resilience to noise and its ability to avoid overestimating cell diversity make it particularly suited for applications involving heterogeneous datasets, such as mapping human disease signatures to model organisms or analyzing archival datasets with incomplete annotations. These attributes underscore the potential of WISpR to bridge gaps in spatial transcriptomics and scRNA-Seq integration, providing reliable insights into tissue heterogeneity and cellular interactions.

The broader applicability of WISpR extends to various biological contexts, including the analysis of tissues with mixed origins or unmatched conditions. Its ability to handle challenging scenarios with mislabeled or absent cell types reinforces its utility in uncovering complex tissue architectures and identifying disease-specific cellular signatures. These findings highlight the robustness and reliability of WISpR as a tool for high-resolution mapping of cellular compositions in diverse datasets, advancing our understanding of tissue organization and disease mechanisms.

#### 2.3.2 Developing Human Embryonic Heart

Previous studies extensively analyzed the human embryonic heart at 6.5 postconceptional weeks (PCW), demonstrating the utility of datasets from this stage for high-resolution tissue mapping through integration and deconvolution model[6, 28, 30, 48, 49]. The original study included spatial transcriptomics, scRNA-Seq, and in situ sequencing analysis of developing human embryonic heart tissue at three stages (4.5-5.0 PCW, 6.5-7.0 PCW and 9.0 PCW), in order to understand cell-type interactions in human organogenesis and obtain a 3D representation of heart development. The objective was to determine the interaction of cell types involved in human organogenesis[37] and to acquire a three-dimensional representation of the development of the human heart topology.

For this study, scRNA-Seq data from 6.5–7.0 PCW human embryonic heart tissue[37] served as the reference, encompassing 15 cell types from 3,717 cells and 15,323 genes. Spatial transcriptomics data, comprising 1,515 capture spots from the same developmental stage, were deconvoluted to spatially localize these 15 cell types. Fig.5 highlights the mapping performance for two cell types: ventricular cardiomyocytes (Fig.5**a**–5**f**) and fibroblast-like cells associated with vascular development in the outflow tract (Fig. 5**g**– 5**l**). Comprehensive predictions for all 15 cell types are provided in Figures S8–S16.

**Fig. 5.**
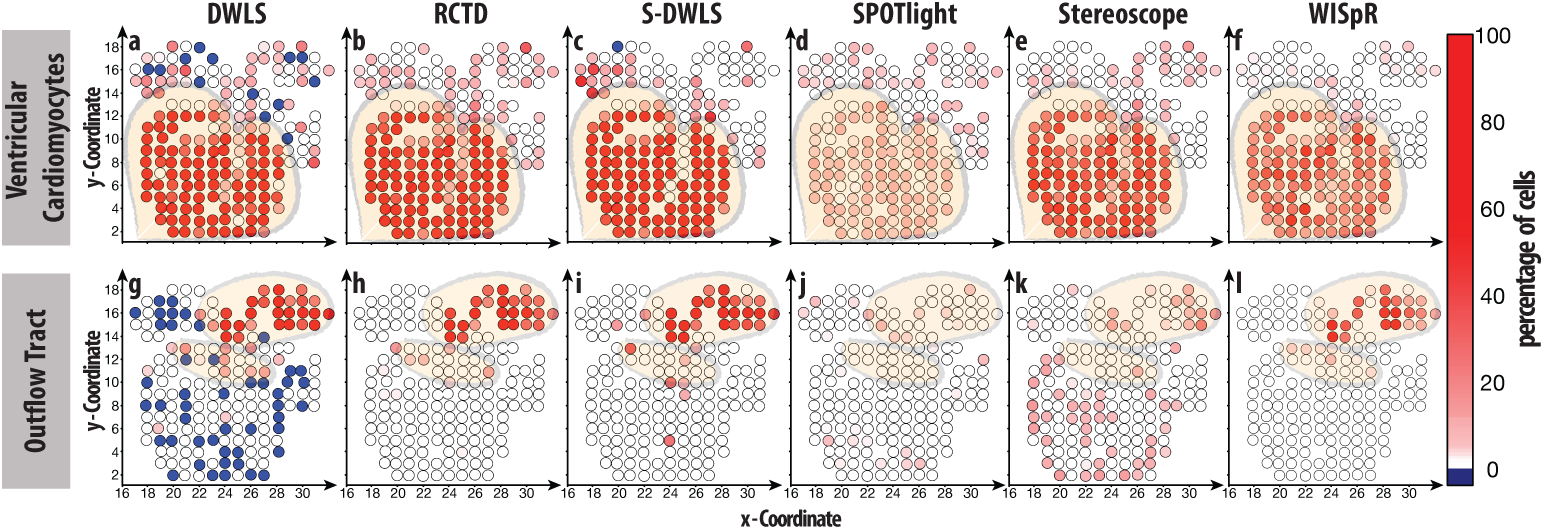
Predictive performance evaluation on real heart data with selected cell types. Top: Predictions for ventricular cardiomyocytes. Bottom: Predictions for outflow tract cells. Blue denotes unrealistic negative predictive coefficients, while the red gradient represents the cell type percentages in the corresponding capture spots. **a** and **g**, DWLS predictions reveal numerous spots with unrealistic negative coefficients. Biologically incorrect positive coefficients are also evident, particularly around the atria and outflow tract, especially for ventricular cardiomyocyte cell prediction. **b** and **h**, RCTD shows a high abundance of false-positive predictions for ventricular cardiomyocyte cells. For the outflow tract, four spots represent false-positive predictions at a low percentage. **c** and **i**, S-DWLS exhibits improved predictions; however, false-positive predictions persist for both ventricular cardiomyocytes and the outflow tract. One spot shows a negative coefficient in **c**. **d** and **j**, SPOTlight predictions indicate overpredicted capture spots in the corresponding tissue. **e** and **k**, Stereoscope also overpredicts cells belonging to ventricular cardiomyocytes. Moreover, cells for the outflow tract are incorrectly predicted in the epicardial zone of the heart. **f** and **l**, WISpR predictions accurately emphasize abundant ventricular cardiomyocytes and outflow tract cells in associated spots.

Although previous efforts made significant progress in annotating cell types and visualizing their spatial distribution, critical limitations in deconvolution accuracy remained, particularly in accounting for the biological sparsity inherent in spatial data. For example, DWLS, RCTD, and Stereoscope frequently produced biologically implausible predictions, such as negative cell-type proportions or overextended spatial assignments. This was particularly evident in tissues with high heterogeneity or low-abundance cell types, such as fibroblast-like cells in the outflow tract or ventricular cardiomyocytes. Models like SPOTlight and DWLS were less effective in resolving the spatial distribution of rare or closely related cell types, often blending predictions across biologically distinct zones. This reduced their ability to map precise cellular interactions critical for understanding heart development. WISpR was specifically designed to address these gaps by introducing biological sparsity constraints and spot-specific optimization, which improve its robustness to noise, mismatched datasets, and rare cell types. By incorporating these features, WISpR offers improved resolution of sparse data and high sensitivity to rare cell types.(Fig. 5 and Figures S6–S14)

Ventricular cardiomyocytes, specialized muscle cells critical to heart function, are localized in the ventricular regions. Immunohistochemistry (IHC) staining with the TNNT2 marker in earlier studies revealed ventricular separation and septum formation as early as 4.5–5.0 PCW[37]. Spatial predictions by various models (DWLS, RCTD, S-DWLS, SPOTlight, Stereoscope, and WISpR) are shown in Fig. 5**a**– 5**f**. DWLS and S-DWLS produced slightly negative predictions (blue) in some regions, indicating biologically implausible outcomes (Fig. 5**a**, 5**c**). RCTD, SPOTlight, and Stereoscope identified ventricular cardiomyocytes in both ventricles and atrial regions but failed to capture their precise spatial distribution or abundance (Fig. 5**b**, 5**d**, 5**e**). WISpR provided the most accurate predictions, correctly localizing ventricular cardiomyocytes and reflecting biologically relevant proportions in each capture spot (Fig. 5**f**).

Advanced vascular development, including the formation of the aorta and pulmonary artery, is evident in the 6.5 PCW heart. Smooth muscle actin (ACTA2) staining in earlier studies highlighted the contribution of fibroblast-like cells to these structures[37]. The deconvolution results for these cells are shown in Fig. 5**g**– 5**l**. DWLS predictions included numerous slightly negative values, further underscoring its limitations in accurately modeling spatial distributions (Fig. 5**g**). RCTD, S-DWLS, and Stereoscope provided improved predictions for the outflow tract compared to ventricular cardiomyocytes. However, their results included an increased number of false-positive capture spots, erroneously extending the predictions to the ventricular regions and the septum (Fig. 5**h**, 5**i**, 5**k**). SPOTlight produced nearly random predictions with sparse biological relevance, reinforcing its unreliability for this application (Fig. 5**j**). WISpR delivered superior results, accurately localizing fibroblast-like cells within biologically relevant regions of the outflow tract with minimal errors (Fig. 5**l**). WISpR consistently outperformed competing methods, providing biologically coherent predictions across different cell types and regions, as validated by spatial stainings of TNNT2 and ACTA2[37]. By minimizing false-positive rates and accurately resolving cell-type proportions, WISpR demonstrated its robustness and precision in deconvolution tasks, particularly for complex tissues such as the developing human heart. These findings underscore WISpR’s ability to integrate spatial and scRNA-Seq datasets effectively, advancing our understanding of cellular organization during organogenesis.

By addressing the challenges with sparse and noisy data and limited sensitivity to rare cell types, WISpR fills a key gap in spatial transcriptomics deconvolution. Not only does it improve the accuracy of cell-type localization in complex tissues such as the developing human heart, it also enables robust integration of spatial and scRNA-Seq data across diverse experimental conditions. This capability is pivotal for advancing our understanding of tissue heterogeneity and uncovering previously hidden cellular interactions during organogenesis.

### 2.4 Applications: WISpR at work

#### 2.4.1 Mouse Brain Cellular Maps

The heterogeneity of the brain, comprising various types of cells such as neurons or astrocytes, underpins its complex functions. The mouse brain section revealed regional clusters (k-means, k = 10) of the Space Ranger program (10X Genomics)) -CTX+AMYG (violet), HY+cerebral peduncle (blue), olfactory sensory neurons (yellow), ventral posterolateral TH (dark green), Ndnf+ neurons (cyan), fiber tracts (magenta), HPF and DG (brown), Npy+ neurons (light green), ventricles (white) and stria terminalis (grey)-each is illustrated in (Fig. 6**a**, 6**b**). *Npy+* neurons (involved in physiological modulation[50–53]) were enriched in the cortical and hippocampus regions (light green), while *Ndnf+* neurons (L1 interneurons[54–56]) were prominent in cortical, HPF[57] and AMYG[58] (cyan).

**Fig. 6.**
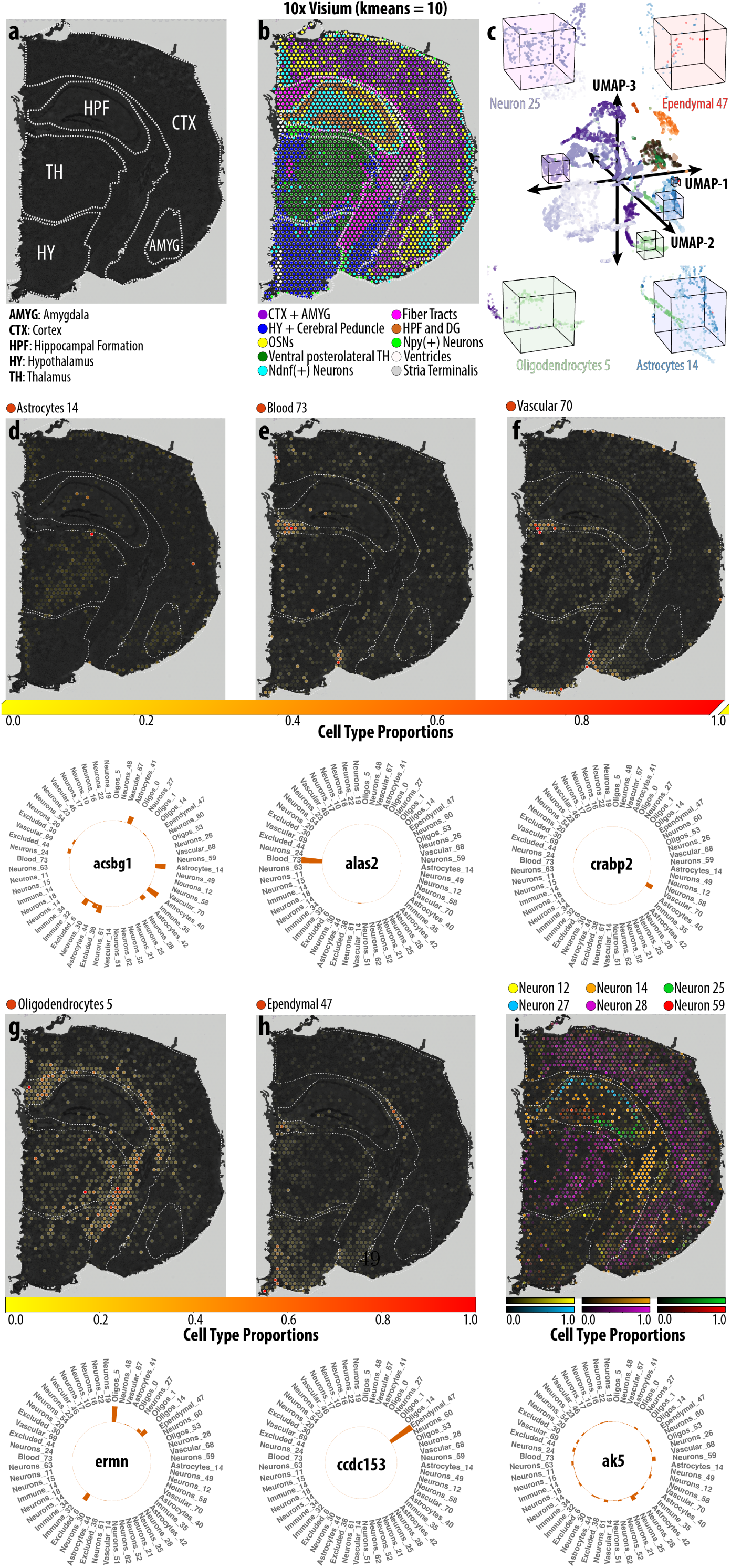
Precise spatial delineation of brain cell types in adult mouse brain. **a**, Illustration of the right hemisphere coronal section of an adult mouse brain, outlining major brain regions with dashed lines. **b**, Identification of 10 spatial clusters, extracted from the 10X Genomics database, representing CTX+AMYG (violet), HY+cerebral peduncle (blue), OSNs (yellow), ventral posterolateral TH (dark green), Ndnf+ neurons (cyan), fiber tracts (magenta), HPF and DG (brown), Npy+ neurons (light green), ventricles (white), and stria terminalis (grey). **c**, 3D UMAP illustration of scRNA-Seq [69], representing major cell types with distinct colors and cellular subtypes as shades of the major color. For simplicity, Neuron 25 (violet), Ependymal 47 (orange), Oligodendrocytes 5 (light green), and Astrocytes 14 (light blue) are depicted. **d-i** Spatial localization of some selected cell types and their selected marker genes are given below. **d**, Spatial localization of Astrocytes 14 on AMYG, CA1, CA3, DG, and outer layers of CTX, with expression levels of a critical astrocyte-specific DEG, *Acsbg1*. **e** and **f**, Coinciding spatial locations of Blood 73 cells and Vascular 70 cells, respectively, on the third ventricle, the ventral intersection point of the cerebral peduncle and the molecular layer of the DG. The expression of *Alas2* and *Crabp2*, which are an erythrocyte and a fibroblast marker, respectively, are enriched. **g**, Cells of the Oligodendrocytes 5 localized in the corpus callosum, fimbria, stria terminalis, and cerebral peduncle, enriched in *Ermn* **k** (left), specifically expressed in myelinating oligodendrocytes. **h**, Ependymal 47 cells predominantly located in the lateral and third ventricles expressing *Ccdc153*, an ependymal cell marker gene. **i**, Representation of six distinctly located neuronal subtypes, Neuron 12 (yellow), Neuron 14 (orange), Neuron 25 (green), Neuron 27 (blue), Neuron 28 (magenta), and Neuron 59 (red). A brain-specific *Ak5*, is differentially expressed within Neuron Cluster 25 but also found in other neuronal cell types and a few excluded cells, indicating their possible neuronal characteristics.

AMYG, involved in the processing of fear and pleasure emotions, interacts with CTX, which contributes to decision making and social behavior[59, 60]. Olfactory sensory neurons (yellow) form spatial clusters in AMYG, mediating sensory input and feedback mechanisms[61]. Cluster analysis highlights complex neural circuits, including the HY (dark blue), the TH (dark green), and HPF (brown), which are linked to diverse functions such as arousal, pain modulation, food aversion, and neurogenesis[62–66]. Key regions such as the paraventricular nucleus (PVT), the lateral habenula, and the cerebral peduncle underscore inter-connectivity within this neural network[67]. The third and lateral ventricles (white) are associated with cerebrospinal fluid generation, while the terminal stria (gray) regulates autonomic and neuroendocrine responses[68]. The preprocessing of scRNA-Seq data[47, 69] involved aggregating metadata from the “Class” and “Clusters” columns to assign cell-type labels. Subcluster attributes were then assigned to the predominant cell types, refining the categorization of the reference cell types. Rigorous cell-type filtering was performed that adhered to the criteria outlined in the study by Andersson *et al.* (2020)[28] was conducted, selecting cell types with a population size ranging from a minimum of 25 to a maximum of 250 cells. The final reference scRNA-Seq data comprises 56 cell types, including 5 Astrocyte, 1 Blood, 1 Ependymal, 4 Excluded, 4 Immune, 30 Neuronal, 5 Oligodendrocytes, and 6 Vascular cell types, totaling 8,449 cells. Although predominant cell populations clustered clearly on the UMAP plot (highlighting localized differences; (Movie S1)), cellular subtypes showed similarities in gene expression profiles, highlighting heterogeneity between cell types (Fig. 1**c**, Fig. 6**c**, Movie S2 and Fig. S21). Accurate deconvolution of these cell types is crucial for mapping spatial distributions and providing insight into neural circuitry and region-specific functions of the brain. In addition, it reveals unique spatial gene expression profiles for each cell type, a critical step in understanding regional functionality. Accordingly, WISpR deconvolution method was employed to determine the precise spatial locations of mouse brain cells, using coronal sections of adult mouse hippocampal tissue in conjunction with reference scRNA-Seq data[47, 69].

The spatial transcriptomics dataset, which contains a coronal section of an adult mouse brain, includes 2,698 capture spots[70], each 55 µm in diameter. To illustrate the predictive outcomes of WISpR, all class and cluster pairings within mouse brain Visium datasets (10X Visium array) were examined (Figure S17–S18). Biologically and regionally distinct cells, astrocytes (Cluster 14), oligodendrocytes (Cluster 5), ependymal cells (Cluster 47), spatially distinct neurons (Clusters 12, 14, 25, 27, 28, and 59), and vascular cells (Clusters 70), were selected for representation in Fig. 6**d**– 6**i**. The selected DEG sets (see the Materials and Methods section for DEG analysis) were further confirmed by conducting enrichment analysis for these six cell types via the Molecular Signatures Database (MSigDB). This enrichment was determined using the markers of the curated cell types present in 233 set of gene identified from single cell sequencing studies carried out on mouse tissue[71] (msigdb.v7.5.1 mm, Category = M8) (Fig. S22). Individual capture spots represent cell-type proportions displayed in varying color shades (Fig. 6**d**– 6**i**). Alongside the predicted cell-type locations, an example of significant DEGs is presented for each cell type whose expression is characteristic of the corresponding cell subtype or the predominant type.

Astrocytes play an important role in structural support, neurotransmitter regulation, and plasticity[72, 73]. Astrocytes (Cluster 14) were predicted to be predominantly localized in the TH and AMYG regions, with a lower representation in CTX, CA1, CA3, and DG of HPF (Fig. 6**d**). The enrichment analysis of cell types of the astrocyte scRNA-Seq cluster (Cluster 14) produced the “large group of glial cells” and “astrocytes”, underscoring the specificity of the DEG set (Fig. S22**a**). In particular, increased expression of acyl-CoA synthetase bubblegum family member 1 (*Acsbg1*), an astrocyte marker, was detected in astrocyte (Cluster 14), coexisting with its expression in a subset of other astrocyte clusters (Clusters 40 and 41).

WISpR’s blood (Cluster 73) and vascular (Cluster 70) cell prediction revealed their co-localizations in key brain regions such as the third ventricle, the ventral intersection of the cerebral peduncle and dentate gyrus (Fig. 6**e**, 6**f**). The enrichment analysis of blood cell types (Cluster 73) revealed its specificity as an erythrocyte (Fig. S22**b**), while the vascular cell types (Cluster 70) emerged as the predominant fibrous cell type, particularly characterized by collagen-rich connective tissue (Fig. S22**e**). Aminolevulinic acid synthase 2 (*Alas2*), an erythroid-specific marker gene, exhibited high expression in Blood Cluster 73, further confirming the findings of the cell-type enrichment analysis. Cellular retinoic acid binding protein II (*Crabp2*), a fibroblast marker specifically expressed in arachnoid and dura fibroblasts in the brain[74], was significantly enriched in vascular Cluster 70, emphasizing the distinct cellular identities of these clusters, as revealed by cell enrichment analysis.

In the scRNA-Seq reference dataset, oligodendrocytes appear as five different cell subtypes with similar gene expression profiles (Fig. 6**c**, S21). The enrichment analysis of cell types highlights Cluster 5 as it consists of oligodendrocyte cells, specifically identified as prostaglandin D2 synthase (*Ptgds*) positive oligodendrocyte precursor cells (Fig. S22**d**). Notably, *Ptgds* ranks among the most abundant transcripts that encode secreted proteins within oligodendrocyte lineage cells[75]. Furthermore, Ermin (*Ermn*), a myelinating oligodendrocyte-specific protein[76], exhibits high expression levels in Oligodendrocyte Cluster 5. This expression pattern aligns with WISpR predictions, localizing this cell type in the corpus callosum, fimbria, stria terminalis, and cerebral peduncle[76, 77] (Fig. 6**g**).

A unique ependymal cell type designated as Cluster 47 was identified as ciliated epithelial glial cells derived from radial glia positioned along the surfaces of the brain ventricles and the spinal canal, as detailed by Macdonald *et al.*. (2021)[78]. The enrichment analysis of cell types supports that the set of genes associated with Cluster 47 exhibits significant enrichment in ciliated glial cells (Fig. S22**c**), and the ependymal cell marker gene *Ccdc153* [79] is specifically expressed within these cell types. In addition to these, WISpR predominantly indicates the location of ependymal cells within the lateral ventricle and the third ventricle, thus attesting to the precision of the localization predictions (Fig. 6**h**).

The gene expression patterns of neurons in Fig. 6**c** exhibit significant similarity, highlighting the presence of cellular heterogeneity (Fig. S21). Nevertheless, WISpR effectively unravels these complexities, offering precise predictions that highlight the distinct spatial distributions of neuronal subtypes within the brain. Among the 30 identified subtypes, Fig. 6**i** portrays six neuronal subtypes. In this context, a predominant location of Neuron Cluster 12 is demonstrated in the posterior parietal, primary somatosensory, and dorsal auditory regions of the CTX, along with the inferior and superior areas of the fiber tracts. Furthermore, cells in neuron (Cluster 14) are predominantly clustered in the stria terminalis. Neuronal Clusters 25, 27, and 59 are clustered within HPF, specifically located in CA3, CA1, and DG, respectively. Finally, Neuron Cluster 28 emerges as a prominent neuronal subtype, with cells predominantly observed in both CTX and TH (Fig. 6**i**), suggesting a functional connectivity across these regions. Upon examination of Neuron Cluster 25, the cell-type enrichment analysis reveals the representation of visceral and excitatory neuron characteristics within this cluster, where these neuronal features are observed in the neurons located in HPF (Fig. S22**f**). Furthermore, adenylate kinase 5 (*Ak5*), a brain-specific marker, is noticeably expressed within Neuron Cluster 25, along with other neuronal subtypes, including Neuron Clusters 59, 60, and 61. By providing precise neuronal maps, WISpR enables insights into brain organization, connectivity, and the cellular basis of neural function and disease.

Using WISpR to integrate spatial transcriptomics with scRNA-Seq data, this study provides a comprehensive high-resolution atlas of cellular heterogeneity and spatial organization in the adult mouse brain. WISpR accurately mapped distinct astrocytes, oligodendrocytes, neurons, blood, vascular, and ependymal cell populations, revealing their spatial distributions and functional annotations. By delineating their region-specific functions, such as the participation of astrocytes in neural plasticity, the contribution of oligodendrocytes to myelination, the subtype-specific connectivity of neurons, and the structural and metabolic functions of vascular cells, WISpR uncovers critical insights into intricate neural circuitry and brain functionality. These findings highlight the potential of WISpR as a transformative tool to advance our understanding of brain organization and support targeted investigations into neural health and disease.

#### 2.4.2 Cancer Origins in Human Breast Cancer

Cancer fundamentally alters cellular characteristics, gene expression profiles, and physiological features[80], driving the need for bottom-up genetic approaches that underpin modern precision oncology[81]. Transcriptomics has become central to cancer research, providing comprehensive insights into the intricate gene expression patterns of cancer cells and their microenvironment[81, 82].Comprehensive transcriptome analysis deciphers the molecular mechanisms driving cancer initiation, progression[83, 84], and therapeutic response[85], where profiling reveals potential diagnostic and prognostic biomarkers, therapeutic targets, and uncovers the molecular heterogeneity of tumors, revealing intricate cancer landscapes[84, 86, 87], offering a holistic view of cell dynamics. Therefore, Deconvoluting cellular properties from spatial cancer data clarifies tumor-stroma-immune interactions, and enables precise estimation of cell-type co-localizations that aligns spatial data with observed expression patterns.

This study used six primary human breast cancer samples—four triple negative breast cancer (TNBC) and two positive estrogen receptors (ER +)—from publicly available data repositories[87, 88]. Spatial datasets were generated using the 10X Visium assay and preprocessed using the Space Ranger software v 1.1.0 (10X Genomics) protocol.

The human breast cancer scRNA-Seq dataset was derived from the three distinct breast cancer subtypes, namely TNBC, human epidermal growth factor receptor 2 (HER2 +) and ER +, through analysis of 11 ER +, 5 HER2 + and 10 primary tumor samples from TNBC[87]. Initially, patient subtyping used the PAM50 method[89, 90], adapted for single-cell data to create “pseudobulk” profiles per tumor, employing the PAM50 centroid predictor. This process also involved pairing tumor cells and analyzing DEGs to identify four distinct gene sets, termed “SCSubtypes” [87], to define single cell-derived molecular subtypes, which were used for external validation of WISpR’s cell-type predictions. These scRNA-Seq datasets consist of 100,064 cells (TNBC: 42,512 cells,(HER2+): 19,311 cells, and ER+: 38,241 cells) and 29,733 genes. This dataset encompasses nine major cell types, including endothelial, cancer-associated fibroblasts (CAF), perivascular-like (PVL), B-cells, T-cells, myeloids, normal epithelial cells, plasmablasts, and cancer epithelial cells, which were further clustered into 29 minor cellular subtypes. The scRNA-Seq cancer types were selected to align with the spatial transcriptomics data by filtering the cellular information to include only the cancer types identified in the spatial dataset. Additionally, subsetting and aligning common genes in both datasets, as illustrated in Fig. 3**b**, 3**d**, produced datasets ready for deconvolution.

WISpR’s spot-specific optimization addresses the challenges of subtype overlaps and rare cell detection by integrating scRNA-Seq data with spatial transcriptomics to achieve biologically coherent deconvolution. During the deconvolution process, optimized parameters for Visium datasets were used, with the spot-specific penalty parameter (*α*) set within the range [0.0–0.3] and the spot-specific thresholding parameter (*τ*) within the range [0.001–0.03]. The six spatial transcriptomics cancer samples were deconvolved using WISpR, with the results for two samples shown in Fig.7 and the remaining results provided in Figures S23–S28.

**Fig. 7.**
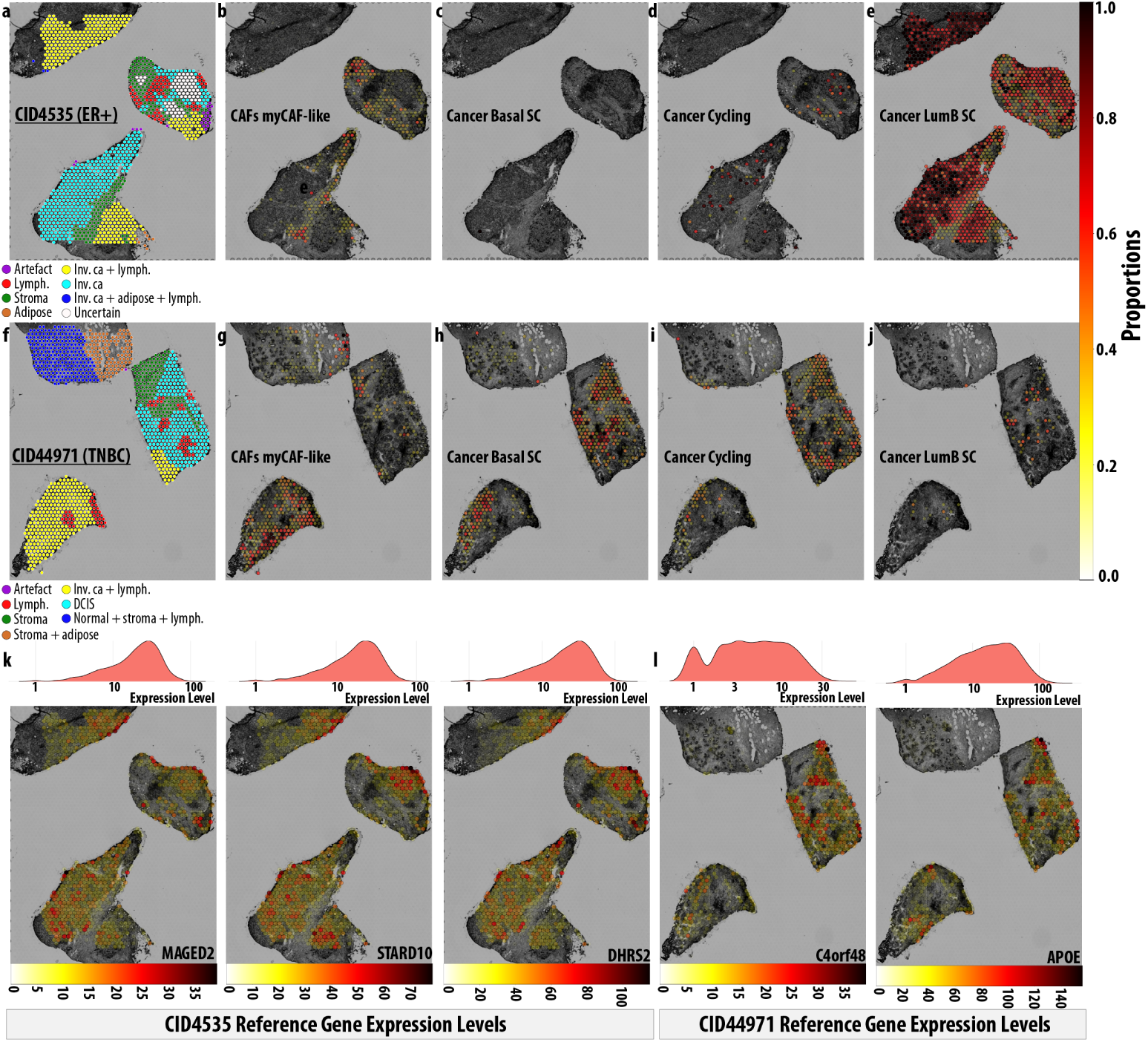
Spatial characterization of cancer-associated cell types in human breast cancer malignancies. **a**, 8 clusters of ER+ breast cancer (Patient ID: CID4535). Invasive cancer-affected tissues were given by shades of cyan, yellow (cancer within lymphatic tissues), and blue (cancer in adipose and lymphatic tissues). Notably, stromal regions (dark-green) and lymphocytes (red) formed distinct clusters without cancerous involvement. Adipose tissue (brown) exhibited minimal representation within the dataset. A cluster labeled as ‘Uncertain’ (white) was identified, alongside sparse occurrences of artifact spots (magenta) [87]. **b**, Localization of CAFs myCAF-like cells, with their abundance proportions in the stroma (less abundance:white, complete abundance:black). **c**, Basal cells were not detected in ER+ tissue sample. **d**, Cycling cells were localized in the “Uncertain” zone. **e**, LumB SCs were highly concentrated in zones annotated as invasive cancer while absent in stromal and lymphatic zones. **f**, 7 clusters of TNBC breast cancer (Patient ID: CID44971). Invasive cancer and lymphatic zones (yellow), and DCIS (cyan). Normal tissue, including lymphocytes and stroma, were represented in blue, with stroma+adipose tissue in brown. Stromal regions were clustered in dark-green. Healthy lymphocytes were distributed across both invasive cancer and DCIS regions but distinctly clustered in red. A few magenta spots were artifacts [87]. **g**, CAFs myCAF-like cells were predominantly predicted in the invasive cancer and stromal+adipose tissue, representing their cancer-prone characteristics. **h**, Basal SCs and **i**, cycling cells were confined on the DCIS and a bounded region of the invasive cancer spatial cluster, representing similar gene expression patterns in distinct zones at high resolution. **j**, The DCIS had the capacity to diversify its cancer microenvironment based on the existence of LumB SCs. **k**, Expression profiles of representative genes, namely *MAGED2*, *STARD10*, and *DHRS2*, for TNBC, and **l** *C4orf48* and *APOE*, for ER+ cancer, identified in “SCSubtypes” algorithm [87]. (Top) The ridge plot represents the distribution of the gene expression levels and (bottom) the spatial distribution of the expression of the genes.

Fig. 7 shows the spatial transcriptomics data for the CID4535 and CID44971 patient samples and a subset of their corresponding WISpR-deconvoluted cell types. In CID4535, an ER + cancer sample, the capture spots were grouped into eight zones: invasive cancer areas (yellow, cyan, dark blue), stroma (dark green), lymphoid tissue (red), and noncancerous adipose tissue (brown), which occupied a small portion of the sample (Fig. 7**a**). Additionally, 22 spots were identified as artifacts (magenta). It is evident that this spatial clustering of zones was insufficient to fully represent the underlying cancer identity and prognosis, as three clusters were designated as invasive cancer with distinct expression profiles. Moreover, two of these three invasive cancer zones include cancers in adipose tissue (blue) and lymphocytes (yellow and blue), where the fundamental tissue profiles are not clearly designated. Also, a cluster named “Uncertain” exhibited the limitation of cancer profiling in spatial transcriptomics data. Therefore, a high-resolution mapping is needed to clarify the distribution of cancer within tissue sections.

Fig.7**b**–7**g** demonstrate WISpR’s ability to resolve specific subtypes within the stromal zones. In particular, myofibroblastic CAF cells (CAFs myCAF-like) are predicted to be exclusively enriched in the stromal and adipose tissue regions in both cancer samples, consistent with previous findings[87, 91] (Fig. 7**b**). WISpR deconvolution revealed no basal cancer stem cells in the ER+ sample (CID4535), consistent with their reported prevalence in TNBCs[87] (Fig. 7**c**). Interestingly, WISpR predicted cancer cycling cell positive spots, some expressing genes such as *SCGB2A2*[92], an important biomarker of metastatic breast cancer, in spatial zones indicated as “Uncertain” (Fig. 7**d**), revealing the predictive efficacy of WISpR. WISpR also identified high LumB stem cell positivity in nonstromal regions, confirming the LumB molecular subtype of this cancer (Fig. 7**e**). This level of resolution, unattainable with clustering-based spatial transcriptomics alone[87], highlights the strength of WISpR to refine broad clusters into biologically meaningful sub-populations, offering critical insights into cellular composition and tumor specialization.

Fig. 7**f** depicts a TNBC (patient ID: CID44971). This sample was clustered into seven zones, yellow indicating invasive cancer and lymphocytes, and cyan representing ductal carcinoma in situ (DCIS), a premalignant breast tissue lesion[93], typically surrounded by layers of myoepithelial cells and a basement membrane structure. A distinct group of lymphocytes is observed in areas where DCIS and invasive cancer emerge (red). Stromal tissue is localized at the border of DCIS (green). A tissue section comprises stroma, adipose tissue, lymphocytes, and normal tissue (brown and blue). Only one spot was identified as an artifact (magenta).

WISpR accurately localized myCAF-like CAFs within lymphoid regions of invasive cancer clusters as well as basal cancer stem cells and cycling cancer cells within the confines of DCIS (Fig.7**g**, 7**i**), revealing spatial patterns indicative of differential proliferative activity between DCIS and invasive cancer zones. These findings highlight WISpR’s precision in resolving spatial heterogeneity, enabling detailed mapping of tumor-stroma dynamics. Interestingly, both basal cancer stem cells and cycling cancer cells are detected within the upper left segment of the spatial cluster comprising invasive cancer and lymphocytes, potentially indicating variances in proliferative activity compared to other invasive cancer zones. In clinics, DCIS lesions were classified by immunohistochemical staining for ER, PR, HER2 and related biomarkers[94]. Interestingly, other studies on the expression of the DCIS genes have revealed its intrinsic subtypes, with a significant percentage (20%) that does not match any of these subtypes, indicating a heterogeneous composition of DCIS[95]. Similarly, the TNBC sample CID44971 with DCIS contained Luminal B cancer stem cells within the DCIS areas, indicating heterogeneity and dual tumorigenesis in this sample (Fig. 7**j**).

The basal SC, HER2+ SC, LumA SC and LumB SC biomarkers were identified by Wu *et al.* (2021) through the “SCSubtypes” algorithm [87]. Of the 65 LumB SC markers, 60 were present in CID4535, and of the 89 Basal SC markers, 82 were present in the spatial data of CID44971, where each of their expression levels are given in Figure S29–S31. Among these marker genes, the expression profiles for the 3 representative genes, namely *MAGED2*, *STARD10* and *DHRS2*, for ER+ breast cancer and 2 representative genes, namely *C4orf48* and *APOE*, for TNBC were given in Fig. 7**k**,7**l**, respectively. Interestingly, a significant subset of these genes shows elevated expression levels, with spatial expression patterns aligning closely with the expected localizations of tumor-associated cell types (Figure S32–S36). Specifically, *MAGED2*, *STARD10*, and *DHRS2* show predominant expression in regions initially identified as invasive cancer sites[87], as well as in spots predicted by WISpR to be LumB stem cell territories. Therefore, leveraging WISpR, marker genes such as *MAGED2* and *C4orf48* were localized to specific tumor microenvironments, enabling a spatially resolved understanding of their roles in tumor progression and therapeutic resistance. In particular, these three genes show low or negligible expression in the stromal and lymphatic regions, underscoring their strong association with ER+ breast cancer (Fig. 7**k**). Additionally, *C4orf48* and *APOE* are prominently expressed in DCIS and invasive cancer sites, as well as in spots where WISpR predicts the presence of CAFs myCAF-like and basal stem cell phenotypes (Fig. 7**l**). However, no expression was detected in normal tissues such as stroma, adipose tissue, and lymphatic tissues, strengthening the link between these two genes and invasive breast cancer, including DCIS.

Consequently, WISpR’s advanced deconvolution capabilities provide a paradigm shifting strategy to understand the spatial heterogeneity and molecular complexity of breast cancer. By integrating scRNA-Seq data with spatial transcriptomics, WISpR accurately resolved cellular subtypes, including invasive cancer cells, basal and cycling cancer stem cells, and myCAF-like CAFs, across diverse tumor microenvironments. This study underscores the precision of WISpR in mapping the dynamics of the tumor stroma, identifying spatially distinct cancer subpopulations, and clarifying ambiguous zones such as DCIS. Importantly, external validation of WISpR predictions using spatial expression patterns of established cancer marker genes—such as *MAGED2*, *STARD10*, and *C4orf48* —demonstrated strong alignment with predicted tumor subpopulations and their microenvironmental contexts, reinforcing the reliability of the model. The ability of WISpR to uncover functional subpopulations and their spatial relationships holds significant promise for advancing cancer research, enabling deeper insights into tumor progression, therapeutic resistance, and the development of precision oncology strategies.

## 3 Discussion

Gene expression regulates cellular functions, responses, and developmental processes. Transcriptomics analysis, enabled by next-generation sequencing, profile differential gene expressions, providing information on gene activation or suppression in response to stimuli. This information is pivotal for understanding cellular responses to environmental signals and signaling pathways, discovering disease mechanisms, identifying therapeutic targets, and advancing precision medicine.

Advances in next-generation sequencing have enabled high-resolution scRNA-Seq and spatial transcriptomics datasets, offering profound biological insights when integrated. Extensive research in transcriptomics has led to the development of efficient techniques to understand not only the spatial location of cells but also their spatial communication and interactions. These insights are crucial for understanding the biological morphology and functionality of biological systems. However, transcriptomics deconvolution faces the challenge of cellular coherence and heterogeneity during the identification and quantification of diverse cell types in complex tissues, especially with unmatched datasets where scRNA-Seq and spatial transcriptomics data do not align completely. When datasets mismatched, reference profiles may not accurately represent the cell types in the sample, leading to errors in identification and quantification. Such mismatched datasets are often used to infer the spatial arrangement of cell types when either dataset is unavailable. However, existing deconvolution methods frequently validate models using synthetic datasets derived from scRNA-Seq, predicting cell types that may not fully reflect real data conditions. Consequently, their real data analysis frequently involves the use of mismatched datasets without sufficient testing of the model performance under these conditions.

We introduce WISpR, a robust deconvolution method for integrating scRNA-Seq and spatial transcriptomics data, tailored to both matched and mismatched datasets. Using biological sparsity and spot-specific optimization, WISpR achieves unparalleled precision in resolving spatial cell-type composition, offering a transformative approach to understand tissue architecture. This breakthrough not only advances computational deconvolution but also opens avenues for deeper biological exploration, such as the identification of region-specific cellular interactions in health and disease.

The limitations of widely used deconvolution models (DWLS, RCTD, S-DWLS, SPOTlight, and Stereoscope) in detecting crucial cell types are emphasized, particularly in scenarios involving the stress test for deconvolution of human heart cells within mouse brain tissue and the biological validation test for human brain HPF cells within whole mouse brain tissue. In addition, almost all models generated divergent spatial patterns for each cell type, which complicates the evaluation of their reliability. Notably, only WISpR accurately represents mouse brain data with biologically acceptable signals, reflecting substantial tissue and organismal, as well as regional differences between scRNA-Seq and spatial data. This accurately reflects the attempt to deconvolute human heart cells in this specific scenario.

The benchmarking on semi-synthetic datasets, including four technical scenarios and six blended datasets, WISpR performance was evaluated against the deconvolution models. Across four scenarios representing the matched datasets, WISpR consistently achieved the lowest RMSE values, ranging from 0.032 to 0.071, and the highest F1 scores, reaching up to 0.905, significantly outperforming five widely used models (DWLS, RCTD, S-DWLS, SPOTlight, and Stereoscope). Compared to S-DWLS, the next model that performs best in most cases, WISpR achieved up to 30% lower RMSE and 4% higher F1 scores (Scenario 1), reflecting its superior ability to accurately locate cell types with minimal errors. Against SPOTlight, which had the highest RMSE values (up to 0.272) and the lowest F1 scores (as low as 0.233), WISpR demonstrated a remarkable 86% improvement in RMSE and a 414% improvement in F1 scores.

In matched datasets (Scenario 1 and 2), WISpR excelled with RMSE reductions up to 16% to 86% compared to other models and consistently maintained F1 scores above 0.868. Even in noisy and mismatched datasets (Scenario 3) or misannotation of cell types (Scenario 4), WISpR maintained its robust performance, achieving up to 15% lower RMSE and 4% higher F1 scores than S-DWLS, while surpassing SPOTlight and Stereoscope by margins exceeding 70% in RMSE and 400% in F1 scores.

In blended examples simulating unmatched data types, WISpR demonstrates superior performance. Particularly in *Mix*[50, 50, 0], where cell types from different tissues (heart and brain) of different organisms (human and mouse) are equally blended in scRNA-Seq, WISpR demonstrates up to 33% lower prediction errors compared to the second best algorithm and consistently high F1 scores (≥ 0.68) compared to other models, underscoring its efficacy in distinguishing significant cells. Additionally, S-DWLS outperforms DWLS, RCTD, SPOTlight, and Stereoscope in *Mix*[50, 50, 0], underscoring its effectiveness in scenarios with notable differences in cell types. However, when similar cells, lacking in synthetic spatial data, are intentionally introduced in scRNA-Seq, all models, including S-DWLS, fail to distinguish existing cell types from nonexistent ones. Despite this, WISpR maintains its robust prediction performance, consistently achieving high accuracy levels regardless of the noise present in the blended data.

WISpR demonstrated robust performance on sixteen spatial datasets encompassing various organs and health conditions, including the developing human heart, mouse brain, and human breast cancer. Specifically, when tested in the human heart, predictions for highly abundant ventricular cardiomyocytes and less abundant smooth muscle cells/fibroblast-like cells in the outflow tract were compared with the benchmarks set by DWLS, RCTD, S-DWLS, SPOTlight and Stereoscope. Interestingly, DWLS and S-DWLS occasionally produced slightly negative cell-type predictions in spots with inconclusive biological values, which were unusually small and deemed nonsignificant. SPOTlight’s predictions were unreliable, with low percentages of positive cell-type predictions in marked spots. RCTD, S-DWLS, and Stereoscope successfully predicted cells in the ventricular zone of the heart, but also exhibited high cell percentages in spots located in the atrial segments. However, Stereoscope failed to predict cells in the outflow tract, indicating a weakness in predicting relatively less abundant cell populations. RCTD and S-DWLS predicted cells in the outflow tract with moderately few false positive predictions. These observations align with predictions from synthetic data, in which RCTD and S-DWLS exhibited significantly low error rates in matched datasets, with visible false-positive predictions in RCTD.

WISpR was then applied to a mouse brain section to deconvolute cell types from the scRNA-Seq data. This dataset comprised 8 cell types consisting of a total of 56 cell subtypes, for which subtype annotations were not originally provided. WISpR accurately reconstructed the cellular landscape of the mouse brain, resolving gene expression correlations between subtypes and mapping cell types to biologically meaningful locations. This precision revealed unannotated subtypes and their spatial context, paving the way for studies on cellular communication and functional roles. WISpR’s mapping of astrocytes, vascular cells, oligodendrocytes, and neurons in the mouse brain was validated by GSEA, confirming their spatial and functional relevance. These findings emphasize the ability of WISpR to clarify cellular subtypes and their interactions, paving the way for further studies on spatial communication and functional profiles.

In six human ER+ and TNBC samples, WISpR generated a high-resolution cancer cell atlas, accurately identifying cancer subtypes and revealing cellular identities in “Uncertain” regions where spatial methods failed to resolve. Notably, WISpR revealed LumB traits in DCIS regions, a novel finding suggesting their potential role in resistance and progression to treatment. Additionally, WISpR localized myCAF-like CAFs in regions mislabeled as noncancerous stroma, underscoring its precision in resolving tumor-stroma interactions. The predictions of WISpR were also validated by the expression levels and localizations of DEGs specific for cancer subtypes, listed by the SCSubtypes algorithm, by comparing the predicted cell type positions and the spatial expression profiles of the provided DEGs. As a result, WISpR’s high-resolution mapping of the cancer microenvironment underscores its transformative potential in precision oncology. By resolving tumor heterogeneity and revealing cell interactions, WISpR lays the groundwork for developing combinatory therapies to overcome resistance and enhance patient care

## 4 Conclusion

Creating a comprehensive cell atlas for healthy and diseased states requires integrating horizontal (same omics) and vertical (different omics) tools. Sparse reconstruction approaches address challenges from cell state variability, technological differences, and dynamic cellular environments by leveraging the inherent sparsity of biological systems to recover cell-type information. Transcriptomics deconvolution methods that incorporate sparse reconstruction must balance maintaining coherence with accounting for biological complexity and variability, a balance crucial for advancing our understanding of tissue composition and function in health and disease. By redefining deconvolution methodologies through sparse regression, WISpR establishes a scalable framework for integrating multi-modal data, paving the way for future innovations in spatial omics. Future work could extend the WISpR application to multimodal data integration, enabling the integration of epigenetic and proteomic layers for a more comprehensive understanding of cellular states. Furthermore, its robustness in noisy datasets suggests potential in clinical settings, such as cancer heterogeneity analysis and prediction of response to treatment.

These results highlight the superior ability of WISpR to uncover spatially distinct cell states, establishing it as an indispensable tool to investigate tissue microenvironments and unravel cellular heterogeneity in complex biological systems. WISpR emerges as a potent instrument for dissecting the biologically sparse makeup of cell types and spatial configurations within intricate tissues. WISpR transcends the limitations of traditional methods, offering not just incremental improvements but a paradigm shift in how spatial transcriptomics data is analyzed. Its potential to uncover new biological insights into health and disease underscores its value as a cornerstone tool in the era of precision medicine.

## 5 Methods

### 5.1 Similarity Analysis

Similarity scores via linear correlations involve calculating the pairwise Pearson’s correlation coefficient between gene expression profiles of all cell types. The Pearson correlation coefficient (*ρ****_x_***,***_y_***) of gene expression profile vectors ***x*** and ***y*** is determined by the formula:

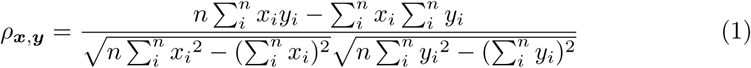

where ***x*** = (*x_i_*), ***y*** = (*y_i_*) ∈ ℕ*^n^* are *n*-dimensional vectors, and *i* = 1*, . . ., n*. The Pearson correlation yields a number from -1 (indicating a perfect negative correlation) to 1 (indicating a perfect positive correlation), with 0 denoting no correlation.

### 5.2 Selection of Differentially Expressed Genes

Differentially Expressed Genes are identified using an algorithm based on the Gini index[45, 46, 96], whose robustness to gene selection has been demonstrated[97]. The Gini index *G* functions as an inequality metric, calculated as the mean absolute divergence among all pairs of data points within the provided dataset. In cases of consistent measurements, the Gini index approaches its minimum value of 0. In heterogeneous cases, the index tends towards the theoretical maximum of 1, indicating the highest inequality[98, 99]. The Gini index for a gene is calculated as:

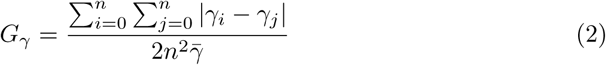

where *i* and *j* represent cell types, *γ* denotes the average expression value observed for a gene in specific cell types, *n* is the number of cell types, and *γ̄* is the mean value of the gene across the entire dataset. Within the algorithm, the average expression values of genes specific to each cell type and the cellular detection rates for the genes in specific cell types are computed. The lower detection rate threshold is set at 0.2, indicating that a gene must be present in at least 20% of cells in a given cell type to be considered a DEG; those below this threshold are excluded from the calculation. Subsequently, the Gini coefficients (GC) are individually calculated for each gene (*γ*), taking into account both the average expression values and the detection rates across all cell types. These computed Gini coefficients for average expression values and detection rates are then transformed into ranking scores, and an integrated score (IS) is computed using all obtained values as follows,

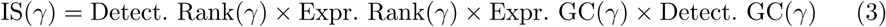

Genes within each cell type *γ_i_* for *i* = 1*, . . ., n*, where *n* is the number of cell types, ranked according to their gene-specific expression and detection ranking, and following Dries *et al.*, (2021)[46] the top 100 genes with the highest values specific to each cell types are used to construct the DEG matrix.

### 5.3 WISpR Model

WISpR effectively obtains a reliable solution to a complex challenge of inverse deconvolution. This problem is formally expressed through a linear equation,

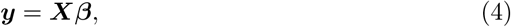

where ***X*** ∈ ℕ*^M×Q^*, ***y*** ∈ ℕ*^M×L^* and ***β*** ∈ ℕ*^Q×L^*. Within this equation, ***y*** is the gene expressions matrix, and each column of ***y*** represents a particular spatial transcriptomic location (that is, a spot), while ***X*** encompasses gene expressions derived from diverse cell types as column vectors, obtained from scRNA-Seq data. The purpose of ***β*** is to capture coefficients that clarify the extent to which cell types within ***X*** influence the spatial transcriptomics data represented by ***y***. Since the total number of gene types exceeds the total number of cell types identified from the scRNA-Seq data (*M > Q*), the deconvolution problem exhibits non-uniqueness in its solutions. In other words, there are infinitely many solutions for Eq. 4. Hence, the challenge lies in discerning the correct solution from the multitude of possibilities.

The fundamental assumption in the deconvolution approach is that one has access to all the necessary scRNA-Seq data for reconstructing spatial transcriptomics. Suppose that spatial transcriptomics data can be represented as linear combinations of a given library ***X*** = {***X***_1_, · · ·, ***X****_Q_*} of scRNA-Seq data, with ***X****_i_* ∈ ℕ*^M^* . For each spot *i*, the objective is to determine a coefficient vector ***β****_i_* = (*β*_1*,i*_*, . . ., β_Q,i_*) that can express the *i*-th component of spatial data ***y****_i_* as a linear combination of ***X***, such that 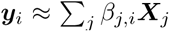. In a matrix representation, we are seeking a coefficient matrix ***β*** so that ***y*** ≈ ***Xβ***, where

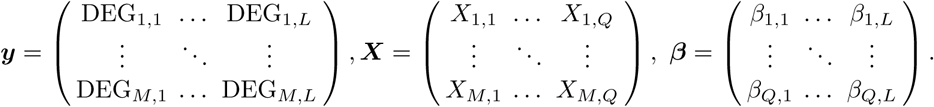

Deconvolution problems are concerned with situations where the number of genes, denoted as *M*, exceeds the number of cell types, represented by *Q*, resulting in an “over-determined” linear equation ***y*** = ***Xβ***. However, considering the inherent sparsity of spatial transcriptomics data, our actual objective is to find a solution for the following minimization problem.

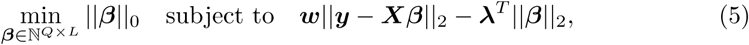

where ||***β***||_0_ := #{(*i, j*) : *β_i,j_≠* 0}, counts the number of non-zero elements in ***β***. In this context, ***λ*** ∈ ℝ is a vector comprising penalty terms, each associated with a specific spot denoted by *i*. These terms play a crucial role in controlling the model overfit by penalizing the coefficients within ***β***. The notation ||.||*_ℓ_* represents a norm used to measure distances in space *L_ℓ_*, i.e., *L*_2_ is the Euclidean distance.

For each spot *i*, we optimize the gene-specific weights indicated as ***w*** = (*w*_1_*, . . ., w_Q_*), with *w_i_* ∈ ℝ^+^. These weights are primarily fine-tuned to mitigate errors that may arise due to the rarity of cell types and gene expressions. To calculate these weights, we employ the multiplicative inverse of the values obtained from ***y****_i_* and use them in sequentially thresholded least squares regressions. It is worth noting that these weights are constrained to be non-negative, effectively ensuring that negative rows are zeroed out, resulting in a weight vector *w_i_* ≥ 0, as exemplified in Algorithm 1.

Throughout each iteration, WISpR computes spot-specific thresholding parameters represented as (*τ_i_*) in Algorithm 1. Coefficients falling below this threshold, indicating nonsignificant coefficients, are eliminated and the regression is conducted sequentially until significant cell-type coefficients are determined.

The sequential thresholding algorithm introduces a specific requirement for ***X*** to guarantee a “unique sparse solution.” This requirement is based on the idea that the columns of ***X*** should exhibit a high degree of orthogonality. This characteristic is of paramount importance, serving as a critical design criterion for our library ***X***, thereby ensuring the uniqueness of our learning objective within a sparse context.

### 5.4 Hyperparameter Tuning

The aim of hyperparameter selection is to optimize model performance and accuracy by adjusting parameters related to complexity and sparsity regularization. In WISpR, three hyperparameter tuning approaches were employed to estimate the optimal three spot-specific parameters, targeting the sparsity parameter (*λ_i_*) to control the sparsity of the model, the weight parameter (*w_i_*) to scale the abundance of cells and gene expression levels, and the thresholding parameter (*τ_i_*) to remove nonsignificant coefficients where *i* = 1*, . . ., L* is the ID of the spot and *L* is the number of spots (Fig. 3).

The parameter *w_i_* is optimized for each individual capture spot using non-negative regression, employing the Limited-memory Broyden, Fletcher, Goldfarb, and Shannon (L-BFGS) algorithm, with temporary *λ* set to 0.1. The multiplicative inverse of this parameter is then used to predict the optimal hyperparameters *λ_i_* and *τ_i_*. GridSearch, using the precalculated *w_i_* parameter, is employed to fine-tune the hyperparameters *λ_i_* and *τ_i_*, with a parameter grid covering a range of values for both *τ_i_* and *λ_i_*, including normalization and intercept point values. The evaluation is based on the Negative

Mean Absolute Error, and cross-validation is conducted with 5 splits and repetitions for robust assessment.

#### Algorithm 1 WISpR algorithm to predict the spatial cell localizations.

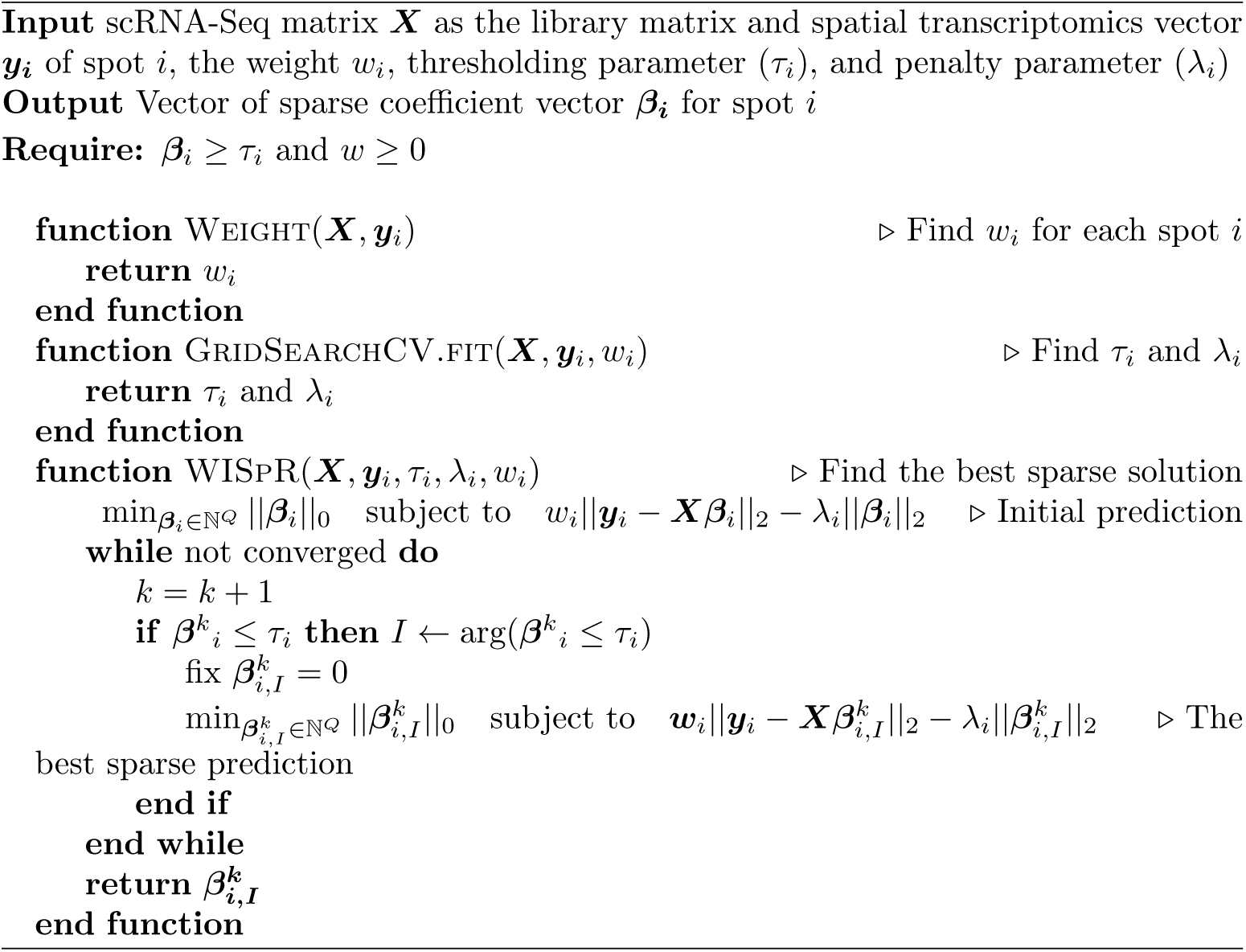

### 5.5 Data-Driven Synthetic Spatial Transcriptomics Reconstruction from scRNA-Seq

Due to the inherent absence of cellular-level gene expression information in real spatial transcriptomics data, evaluating the accuracy of the deconvolution model requires synthetic datasets. These synthetic datasets are expected to mirror the fundamental characteristics of spatial transcriptomics data to simulate real-life deconvolution procedures, encompassing a maximal number of cells and cell types within a synthetic spot. For performance comparisons of deconvolution models, semi-synthetic spatial transcriptomics datasets were prepared using the count matrices and the cell-type annotations given in the scRNA-Seq data based on the methodology explained in Andersson *et al.* (2020)[28].

The synthetic data generation algorithm utilizes scRNA-Seq data, which is represented as a cell × gene matrix where the cell type information for each cell is known. Suppose that the number of available cell types is *N* for the scRNA-Seq data, and the boundaries for the number of cell types (*ct*_min_ and *ct*_max_) and the number of cells (*c*_min_ and *c*_max_) per spot are set. In this work, we choose *ct*_min_ = 1 and *ct*_max_ = 5. The selections for *c*_min_ and *c*_max_ can vary depending on the sequencing technology used.

First, the number of cell types is drawn uniformly (without replacement) from the *N* available cell types within the range of *ct*_min_ and *ct*_max_. Then, the Dirichlet distribution (with a concentration parameter of 1) is employed to probabilistically determine the abundance of cells per cell type in the spot within the selected boundaries of the number of cells (*c*_min_ and *c*_max_). These determined numbers are real numbers, so the relative proportions of the cells are rounded to the nearest integers to maintain a biologically meaningful cell distribution. Finally, the maximum number of cells per cell type is randomly sampled from the available cells. For each sampled cell, the gene expression values are multiplied by a ratio factor (ratio factor =0.1[28]) and added to the spot. Using this algorithm as a foundation, synthetic datasets are prepared to simulate the possible scenarios introduced in the following sections.

#### 5.5.1 Scenarios for Technologies and Boundaries

This section will explore four distinct scenarios related to potential challenges in real RNA sequencing technologies, emphasizing their properties and limitations.

**Scenario 1: Spatial Transcriptomics.** The in-situ barcoding-based spatial transcriptomics technique was introduced in 2016[14]. Since its inception, researchers have rapidly adopted and extensively applied this technique in various studies, including transcriptomics analyses of plant species, developmental studies, cancer research, and investigations into neurodegenerative diseases[100–103].

The spatial transcriptomics slides contain capture spots, each featuring approximately 200 million polydT−oligo-based capture probes. These probes bind to the existing RNAs present in the corresponding tissue position, with each capture probe having a diameter of about 100 µm. The number of cells within a single capture spot can range from 10 to 30, depending on tissue type and location of the section. Thus, it can be regarded as a *bulk RNA* sample on a smaller scale. Our synthetic spatial transcriptomics dataset was created following the same principles as the original data, maintaining a total cell count per spot ranging from 10 to 30 and including approximately 1 to 5 cell types selected from available scRNA-Seq datasets[37]. Three independent synthetic spatial transcriptomics datasets, each containing 500 spots and 15323 genes, were prepared to evaluate the prediction accuracy of the WISpR model.

**Scenario 2: 10X Genomics Visium.** Advancements in in-situ barcoding-based spatial transcriptomics techniques have led to the development of the 10X Visium platform (10X Genomics). This platform allows for spatially resolved gene expression analysis across tissue sections with higher resolution. The reduction in the area of the capture spot from 100 *µ*m to 55 *µ*m has enabled the capture of gene expression profiles from a smaller number of cells per capture spot, with the total number of cells typically ranging from 1 to 10.

To simulate data similar to those generated by the 10X Visium platform, the synthetic dataset for this scenario was created by sampling the total number of cells per capture spot, which varied between 1 and 10. Additionally, the number of cell types per spot was randomly selected to range from 1 to 5. Three independent synthetic spatial transcriptomics datasets were generated to evaluate the prediction accuracy. Each dataset consisted of 500 spots and 15323 genes, ensuring a robust assessment of the WISpR model’s prediction accuracy.

**Scenario 3: Unmatched Cell Types Cases** Comparing the potential cell numbers analyzed by spatial transcriptomics and scRNA-Seq reveals that spatial transcriptomics surpasses scRNA-Seq in both the quantity and variety of cells. This difference can lead to the inclusion of additional cell types in spatial transcriptomics, depending on tissue type, which may not be present in the reference scRNA-Seq data. As a result, these distinct cell types in spatial transcriptomics remain unidentified in the reference data, contributing to noise in the spatial dataset.

To simulate a noisy dataset, synthetic spatial datasets were generated using all 15 cell types in the reference scRNA-Seq data, resulting in three independent datasets, each comprising 500 spots and 15323 genes. Subsequently, the atrial cardiomyocyte cell type ranked as the sixth most populous cell type with 152 cells among the fifteen identified clusters, was excluded from the reference scRNA-Seq sequencing dataset to introduce noise.

**Scenario 4: Mislabeled Cell Types.** Another possible and realistic scenario involves cases where cell types are mislabeled in the metadata.

To simulate this scenario, we initially generated synthetic data following the approach outlined in Scenario 1, utilizing 15 cell types. We produced three distinct synthetic datasets, each consisting of 500 spots and 15323 genes. Subsequently, all 462 cells in cell type 2 were relabeled as cell type 3 in the modified scRNA-Seq metadata. Consequently, the metadata included 14 distinct cell-type labels designated for use in the WISpR deconvolution process.

#### 5.5.2 Blended Data

The accuracy of the deconvolution algorithms in handling unmatched cell types was assessed by controlled simulations. Initially, a synthetic spatial dataset with 1000 spots was generated, incorporating 2923 mouse brain cells from various cell types: Astrocytes 14, Astrocytes 40, Astrocytes 41, Blood 73, Ependymal 47, Excluded 38, Immune 32, Immune 34, Immune 35, Neurons 25, Neurons 26, Neurons 27, Neurons 63, Oligos 0, Oligos 1, Oligos 14, Vascular 14, Vascular 67, Vascular 69.

The cells and cell types utilized in the synthetic data were consistent with those found in the scRNA-Seq dataset, representing 50% of the total scRNA-Seq dataset. To introduce unmatched cells, the remaining 50% consisted of a random selection of cells from the entire human developing heart dataset and non-overlapping cell types from the mouse brain. This selection included Astrocytes 42, Oligos 5, Neurons 48, Oligos 53, Vascular 68, Neurons 12, Neurons 14, Neurons 21, Neurons 52, Neurons 51, Neurons 18, Neurons 15, Neurons 11, Neurons 24, and Neurons 23 as outlined in Table 1.

**Table 1.** Generated blended data using mouse brain cells and human embryonic heart cells, labeled as *Mix*[*a, b, c*], indicates the proportion of mouse brain cells found in both the reference scRNA-Seq and synthetic spatial datasets (*a*), human embryonic heart data (*b*), and mouse brain cells absent in the synthetic spatial dataset within the reference data (*c*).

| Condition | ST(+) | Human Heart | ST(-) |
| --- | --- | --- | --- |
| $Mix[50, 50, 0]$ | 2923 (50%) | 2923 (50%) | 0 (0%) |
| $Mix[50, 40, 10]$ | 2923 (50%) | 2338 (40%) | 585 (10%) |
| $Mix[50, 30, 20]$ | 2923 (50%) | 1754 (30%) | 1169 (20%) |
| $Mix[50, 20, 30]$ | 2923 (50%) | 1169 (20%) | 1754 (30%) |
| $Mix[50, 10, 40]$ | 2923 (50%) | 585 (10%) | 2338 (40%) |
| $Mix[50, 0, 50]$ | 2923 (50%) | 0 (0%) | 2923 (50%) |

### 5.6 State-of-the-art Deconvolution Approaches

To evaluate the predictive performance of the WISpR method, each synthetically generated spatial transcriptomics scenario and the developing human embryonic heart data were deconvoluted and compared with state-of-the-art techniques, including Dampened Weighted Least Square (DWLS)[27], Spatial DWLS[29], Robust Cell Type Decomposition (RCTD)[26], Stereoscope[28], and SPOTlight[30] models.

#### 5.6.1 DWLS

The Dampened Weighted Least Squares (DWLS)[27] model employs vector decomposition-based optimization to address the deconvolution problem, particularly tailored for bulk RNA-seq datasets. The objective is to solve a linear regression equation ***y*** = ***Sx***, where ***y*** represents the gene expression values of the RNA-Seq data in general. The scRNA-Seq signature genes are predicted using the hurdle model implemented in the MAST R package[104], with an absolute log2 mean fold change >0.5, to identify the best cell types ***x*** based on the scRNA-Seq signature gene matrix ***S*** and gene expression vector ***y***. The scRNA-Seq static gene signature matrix ***S*** encompasses the gene expression vector of selected marker genes for corresponding cell types. A weighted error function is utilized to determine the optimal ***x***, which is particularly crucial when dealing with datasets containing rare cell types to avoid overlooking low-expressed informative genes during predictions. To perform the analysis, the publicly available DWLS source code with default parameters is used from the following data repository: https://github.com/dtsoucas/DWLS.

#### 5.6.2 Spatial DWLS

The Spatial DWLS method[29] integrates a Parametric Analysis of Gene Set Enrichment (PAGE) step prior to the DWLS deconvolution process. Furthermore, the Giotto model[46] is utilized to generate the signature gene matrix. The deconvolution procedure in this study was carried out using the code available on GitHub: https://github.com/rdong08/spatialDWLS_dataset.

#### 5.6.3 RCTD

The Robust Cell Type Deconvolution (RCTD) method[26], which employs a modelbased approach to estimate the relative abundance of cells at capture points. Initially, the average mRNA count per cluster is computed. Then, a Maximum Likelihood Estimation (MLE)-based approach is employed to predict cell types. The code and hyperparameters were selected based on the recommendations provided in the GitHub tutorial: https://github.com/dmcable/spacexr.

#### 5.6.4 Stereoscope

The Stereoscope method[28] utilizes a model-based negative binomial deconvolution approach to predict cell-type abundance in spots and identify differentially expressed genes per cell type. Following the official documentation of Stereoscope, the top 5,000 genes that were expressed highest in single-cell data were selected. The algorithm was run with 50,000 epochs, while the remaining parameters were kept as default, as outlined in the repository: https://github.com/almaan/stereoscope.

#### 5.6.5 SPOTlight

The SPOTlight method[30] combines non-negative matrix factorization (NMF) and non-negative least squares (NNLS) for the analysis of transcriptomic data. It operates in three main steps: First, the scRNA-Seq data matrix is decomposed into lowerdimensional matrices, capturing cell-type-specific gene expression patterns and their presence in each cell, called topics. Initialization involves seeding cell-type-specific gene expression patterns with unique marker genes for each cell type and assigning binary values to topics based on cell-type membership. Secondly, it employs NNLS regression to map the transcriptomic data to the specific patterns of the cell type in topics, generating the distributions of the topic profile for each capture location. (referred to as NMFreg). Lastly, it derives consensus cell-type-specific signatures from topics and determines the weights of each cell type present at capture locations using NNLS. In this study, the default parameters recommended for implementation were followed and applied, as indicated in the associated code repository: https://marcelosua.github.io/SPOTlight/.

### 5.7 Performance Comparison Tools

The performance of the 6 deconvolution models was evaluated by comparing their accuracy in estimating each cell type per spatial capture spot using the generated synthetic spatial datasets.

#### 5.7.1 Root Mean Squared Error

The performance of the method was compared by calculating the *Root Mean Squared Error* (RMSE), When the total number of spots was given as *L*, for each spot *ℓ*,

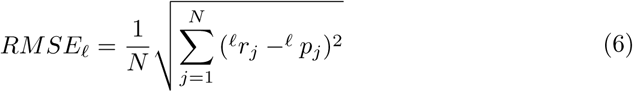

where (*^ℓ^r_j_*) is the ground truth of cell type proportions and (*^ℓ^p_j_*) is the predicted number from the deconvolution models. Here, *j* = 1*, . . ., N* is the index of cell types, and *N* is the total number of cell types.

To statistically analyze the RMSE scores of the deconvolution models, the *Wilcoxon rank-sum test*[105, 106] was employed. This widely recognized statistical hypothesis test is utilized to compare the distributions of two datasets by employing two matched samples. In this work, the built-in test function in the R programming language[107] was utilized with the following parameters:

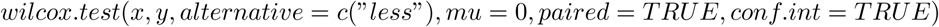

where *x* represents the RMSE values of WISpR at all capture spots, and *y* denotes the RMSE values of the considered alternative methods at all capture spots. The parameter *“alternative = c(“less”)“* is used to statistically analyse whether the RMSE distribution of the WISpR method is closer to zero compared to others.

#### 5.7.2 Total Variational Distance

The difference between false positive predictions and true distributions for each spot *x* for *b* and *c* in *Mix*[*a, b, c*] is calculated as;

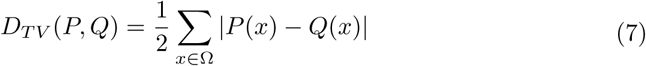

where *P*(*x*) and *Q*(*x*) are the cell-type distributions for prediction and ground truth (zero value), respectively for the Ω number of spots *x*.

#### 5.7.3 F1 Score

*Precision* is a metric that is used to evaluate the performance of the prediction model. Measures the proportion of true positive predictions (correctly predicted positive instances) of all positive predictions that the model makes. In other words, precision measures the accuracy of the model’s positive predictions given as

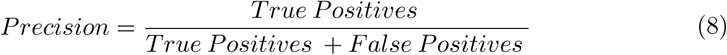

where *T rue Positives* represent the number of correctly predicted positive instances and *False Positives* represent the number of incorrectly predicted positive instances.

*Recall*, also known as the sensitivity rate, quantifies the proportion of true positive predictions out of all actual positive instances. It is calculated using the equation:

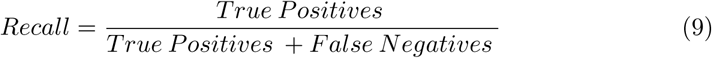

where *False Negatives* represents the number of positive instances incorrectly classified as negative.

*F1 Score*, which combines precision and recall, offers a comprehensive evaluation of the performance of the classification or prediction model. It serves as a balance between precision and recall, which is particularly beneficial for unbalanced data. The F1 Score is calculated using the following formula:

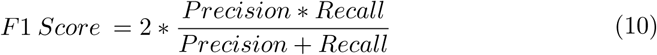

The F1 score ranges from 0 to 1, with 1 being the best possible score. A higher F1 score indicates better model performance in terms of precision and recall.

## 6 Supplementary information

Supplementary Text

Supplementary Figs. S1 to S36

Supplementary Tables S1 and S2

Supplementary Movies S1 and S2

## 7 Acknowledgements

We thank A.O. Argunsah for his helpful comments on mouse brain analysis, and G. Korkmaz for valuable discussions with her on human breast cancer results.

## 8 Declarations

### 8.2 Data and Materials availability

The scRNA-seq and spatial transcriptomics data for human developing heart data can be obtained from https://www.spatialresearch.org/resources-published-datasets/doi-10-1016-j-cell-2019-11-025/. The Visium data for the mouse brain can be obtained from https://support.10xgenomics.com/spatial-gene-expression/datasets/1.0.0/V1_Adult_Mouse_Brain. The scRNA-Seq data for mouse brain L1 hippocampus can be obtained from https://storage.googleapis.com/linnarsson-lab-loom/l1_hippocampus.loom and the cells in each cell type are filtered as cells ≤ 25 and cells ≥ 250. The scRNA-Seq for breast cancer patient data can be downloaded from GSE176078(https://www.ncbi.nlm.nih.gov/geo/query/acc.cgi?acc=GSE176078 and the matching spatial data can be downloaded from https://zenodo.org/record/4739739 [87]

### 8.3 Code availability

The implementation of the WISpR method and its applications are publicly accessible on Zenodo under the DOI: 10.5281/zenodo.11109636 (immediately upon acceptance of the manuscript).

### 8.4 Author contribution

N.S.E. and D.E. developed the WISpR method. D.E. formulated the theoretical approach, while N.S.E. conducted the data analysis and wrote the software. Both N.S.E. and D.E. contributed to the writing, revision, and finalization of the manuscript.

### 8.6 Ethics approval and consent to participate

Not Applicable

### 8.7 Consent for publication

All authors reviewed and approved the final manuscript.

## References

[1] Shah, S., Lubeck, E., Zhou, W., Cai, L.: In situ transcription profiling of single cells reveals spatial organization of cells in the mouse hippocampus. Neuron 92(2), 342–357 (2016)

[2] Menche, J., Sharma, A., Kitsak, M., Ghiassian, S.D., Vidal, M., Loscalzo, J., Barabási, A.-L.: Uncovering disease-disease relationships through the incomplete interactome. Science 347(6224), 1257601 (2015)

[3] Akula, N., Baranova, A., Seto, D., Solka, J., Nalls, M.A., Singleton, A., Ferrucci, L., Tanaka, T., Bandinelli, S., Cho, Y.S., et al.: A network-based approach to prioritize results from genome-wide association studies. PloS one 6(9), 24220 (2011)

[4] Lee, I., Blom, U.M., Wang, P.I., Shim, J.E., Marcotte, E.M.: Prioritizing candidate disease genes by network-based boosting of genome-wide association data. Genome research 21(7), 1109–1121 (2011)

[5] Iossifov, I., Zheng, T., Baron, M., Gilliam, T.C., Rzhetsky, A.: Geneticlinkage mapping of complex hereditary disorders to a whole-genome molecularinteraction network. Genome research 18(7), 1150–1162 (2008)

[6] Rao, A., Barkley, D., França, G.S., Yanai, I.: Exploring tissue architecture using spatial transcriptomics. Nature 596(7871), 211–220 (2021)

[7] Bäckdahl, J., Franzén, L., Massier, L., Li, Q., Jalkanen, J., Gao, H., Andersson, A., Bhalla, N., Thorell, A., Rydén, M., et al.: Spatial mapping reveals human adipocyte subpopulations with distinct sensitivities to insulin. Cell metabolism 33(9), 1869–1882 (2021)

[8] Olaniru, O.E., Kadolsky, U., Kannambath, S., Vaikkinen, H., Fung, K., Dhami, P., Persaud, S.J.: Single-cell transcriptomic and spatial landscapes of the developing human pancreas. Cell Metabolism 35(1), 184–199 (2023)

[9] Melo Ferreira, R., Freije, B.J., Eadon, M.T.: Deconvolution tactics and normalization in renal spatial transcriptomics. Frontiers in Physiology 12, 812947 (2022)

[10] Andersson, A., Larsson, L., Stenbeck, L., Salmén, F., Ehinger, A., Wu, S.Z., Al-Eryani, G., Roden, D., Swarbrick, A., Borg, Å., et al.: Spatial deconvolution of her2-positive breast cancer delineates tumor-associated cell type interactions. Nature communications 12(1), 6012 (2021)

[11] Rájová, J., Davidsson, M., Avallone, M., Hartnor, M., Aldrin-Kirk, P., Cardoso, T., Nolbrant, S., Mollbrink, A., Storm, P., Heuer, A., et al.: Deconvolution of spatial sequencing provides accurate characterization of hesc-derived da transplants in vivo. Molecular Therapy-Methods & Clinical Development 29, 381–394 (2023)

[12] Piwecka, M., Rajewsky, N., Rybak-Wolf, A.: Single-cell and spatial transcriptomics: deciphering brain complexity in health and disease. Nature Reviews Neurology, 1–17 (2023)

[13] Rodriques, S.G., Stickels, R.R., Goeva, A., Martin, C.A., Murray, E., Vanderburg, C.R., Welch, J., Chen, L.M., Chen, F., Macosko, E.Z.: Slide-seq: A scalable technology for measuring genome-wide expression at high spatial resolution. Science 363(6434), 1463–1467 (2019)

[14] Ståhl, P.L., Salmén, F., Vickovic, S., Lundmark, A., Navarro, J.F., Magnusson, J., Giacomello, S., Asp, M., Westholm, J.O., Huss, M., et al.: Visualization and analysis of gene expression in tissue sections by spatial transcriptomics. Science 353(6294), 78–82 (2016)

[15] Kruse, F., Junker, J.P., Van Oudenaarden, A., Bakkers, J.: Tomo-seq: A method to obtain genome-wide expression data with spatial resolution. In: Methods in Cell Biology vol. 135, pp. 299–307. Elsevier, Amsterdam, Netherlands (2016)

[16] Chen, A., Liao, S., Cheng, M., Ma, K., Wu, L., Lai, Y., Qiu, X., Yang, J., Xu, J., Hao, S., et al.: Spatiotemporal transcriptomic atlas of mouse organogenesis using dna nanoball-patterned arrays. Cell 185(10), 1777–1792 (2022)

[17] Fu, X., Sun, L., Chen, J., Dong, R., Lin, Y., Palmiter, R., Lin, S., Gu, L.: Continuous polony gels for tissue mapping with high resolution and rna capture efficiency. bioRxiv (2021)

[18] Cho, C.-S., Xi, J., Si, Y., Park, S.-R., Hsu, J.-E., Kim, M., Jun, G., Kang, H.M., Lee, J.H.: Microscopic examination of spatial transcriptome using seq-scope. Cell 184(13), 3559–3572 (2021)

[19] Hu, Z., Ahmed, A.A., Yau, C.: Cider: an interpretable meta-clustering framework for single-cell rna-seq data integration and evaluation. Genome Biology 22(1), 1–21 (2021)

[20] Brennecke, P., Anders, S., Kim, J.K., Kołodziejczyk, A.A., Zhang, X., Proserpio, V., Baying, B., Benes, V., Teichmann, S.A., Marioni, J.C., et al.: Accounting for technical noise in single-cell rna-seq experiments. Nature methods 10(11), 1093–1095 (2013)

[21] Iacono, G., Massoni-Badosa, R., Heyn, H.: Single-cell transcriptomics unveils gene regulatory network plasticity. Genome biology 20, 1–20 (2019)

[22] Kharchenko, P.V., Silberstein, L., Scadden, D.T.: Bayesian approach to singlecell differential expression analysis. Nature methods 11(7), 740–742 (2014)

[23] L Lun, A.T., Bach, K., Marioni, J.C.: Pooling across cells to normalize single-cell rna sequencing data with many zero counts. Genome biology 17(1), 1–14 (2016)

[24] Grün, D., Kester, L., Van Oudenaarden, A.: Validation of noise models for singlecell transcriptomics. Nature methods 11(6), 637–640 (2014)

[25] Kim, J.K., Kolodziejczyk, A.A., Ilicic, T., Teichmann, S.A., Marioni, J.C.: Characterizing noise structure in single-cell rna-seq distinguishes genuine from technical stochastic allelic expression. Nature communications 6(1), 8687 (2015)

[26] Cable, D.M., Murray, E., Zou, L.S., Goeva, A., Macosko, E.Z., Chen, F., Irizarry, R.A.: Robust decomposition of cell type mixtures in spatial transcriptomics. Nature Biotechnology 40(4), 517–526 (2022)

[27] Tsoucas, D., Dong, R., Chen, H., Zhu, Q., Guo, G., Yuan, G.-C.: Accurate estimation of cell-type composition from gene expression data. Nature communications 10(1), 1–9 (2019)

[28] Andersson, A., Bergenstråhle, J., Asp, M., Bergenstråhle, L., Jurek, A., Fernández Navarro, J., Lundeberg, J.: Single-cell and spatial transcriptomics enables probabilistic inference of cell type topography. Communications biology 3(1), 1–8 (2020)

[29] Dong, R., Yuan, G.-C.: Spatialdwls: accurate deconvolution of spatial transcriptomic data. Genome biology 22(1), 1–10 (2021)

[30] Elosua-Bayes, M., Nieto, P., Mereu, E., Gut, I., Heyn, H.: Spotlight: seeded nmf regression to deconvolute spatial transcriptomics spots with single-cell transcriptomes. Nucleic acids research 49(9), 50–50 (2021)

[31] Kleshchevnikov, V., Shmatko, A., Dann, E., Aivazidis, A., King, H.W., Li, T., Elmentaite, R., Lomakin, A., Kedlian, V., Gayoso, A., et al.: Cell2location maps fine-grained cell types in spatial transcriptomics. Nature biotechnology 40(5), 661–671 (2022)

[32] Geras, A., Darvish Shafighi, S., Domżał, K., Filipiuk, I., Rączkowska, A., Szymczak, P., Toosi, H., Kaczmarek, L., Koperski, Ł., Lagergren, J., et al.: Celloscope: a probabilistic model for marker-gene-driven cell type deconvolution in spatial transcriptomics data. Genome Biology 24(1), 120 (2023)

[33] Montgomery, D.C., Peck, E.A., Vining, G.G.: Introduction to Linear Regression Analysis, 6th edn. John Wiley & Sons, Hoboken, NJ (2021)

[34] Kiselev, V.Y., Kirschner, K., Schaub, M.T., Andrews, T., Yiu, A., Chandra, T., Natarajan, K.N., Reik, W., Barahona, M., Green, A.R., et al.: Sc3: consensus clustering of single-cell rna-seq data. Nature methods 14(5), 483–486 (2017)

[35] Hao, Y., Hao, S., Andersen-Nissen, E., Mauck III, W.M., Zheng, S., Butler, A., Lee, M.J., Wilk, A.J., Darby, C., Zager, M., et al.: Integrated analysis of multimodal single-cell data. Cell 184(13), 3573–3587 (2021)

[36] Guo, M., Wang, H., Potter, S.S., Whitsett, J.A., Xu, Y.: Sincera: a pipeline for single-cell rna-seq profiling analysis. PLoS computational biology 11(11), 1004575 (2015)

[37] Asp, M., Giacomello, S., Larsson, L., Wu, C., Fürth, D., Qian, X., Wärdell, E., Custodio, J., Reimegård, J., Salmén, F., et al.: A spatiotemporal organ-wide gene expression and cell atlas of the developing human heart. Cell 179(7), 1647–1660 (2019)

[38] McInnes, L., Healy, J., Melville, J.: Umap: Uniform manifold approximation and projection for dimension reduction. arXiv preprint arXiv:1802.03426 (2018)

[39] Habib, N., Li, Y., Heidenreich, M., Swiech, L., Avraham-Davidi, I., Trombetta, J.J., Hession, C., Zhang, F., Regev, A.: Div-seq: Single-nucleus rna-seq reveals dynamics of rare adult newborn neurons. Science 353(6302), 925–928 (2016)

[40] Joglekar, A., Prjibelski, A., Mahfouz, A., Collier, P., Lin, S., Schlusche, A.K., Marrocco, J., Williams, S.R., Haase, B., Hayes, A., et al.: A spatially resolved brain region-and cell type-specific isoform atlas of the postnatal mouse brain. Nature Communications 12(1), 463 (2021)

[41] Brunton, S.L., Proctor, J.L., Kutz, J.N.: Discovering governing equations from data by sparse identification of nonlinear dynamical systems. Proceedings of the national academy of sciences 113(15), 3932–3937 (2016)

[42] Satija, R., Farrell, J.A., Gennert, D., Schier, A.F., Regev, A.: Spatial reconstruction of single-cell gene expression data. Nature biotechnology 33(5), 495–502 (2015)

[43] Mallick, H., Chatterjee, S., Chowdhury, S., Chatterjee, S., Rahnavard, A., Hicks, S.C.: Differential expression of single-cell rna-seq data using tweedie models. Statistics in medicine 41(18), 3492–3510 (2022)

[44] Shi, Y., Lee, J.-H., Kang, H., Jiang, H.: A two-part mixed model for differential expression analysis in single-cell high-throughput gene expression data. Genes 13(2), 377 (2022)

[45] Jiang, L., Chen, H., Pinello, L., Yuan, G.-C.: Giniclust: detecting rare cell types from single-cell gene expression data with gini index. Genome biology 17(1), 1–13 (2016)

[46] Dries, R., Zhu, Q., Dong, R., Eng, C.-H.L., Li, H., Liu, K., Fu, Y., Zhao, T., Sarkar, A., Bao, F., et al.: Giotto: a toolbox for integrative analysis and visualization of spatial expression data. Genome biology 22(1), 1–31 (2021)

[47] Cell Scatterplot, Hippocampus. http://loom.linnarssonlab.org/dataset/cells/Mousebrain.org.level1/L1_Hippocampus.loom (accessed: 21.11.2022)

[48] Song, Q., Su, J.: Dstg: deconvoluting spatial transcriptomics data through graphbased artificial intelligence. Briefings in Bioinformatics 22(5), 414 (2021)

[49] Zhuang, X.: Spatially resolved single-cell genomics and transcriptomics by imaging. Nature methods 18(1), 18–22 (2021)

[50] Ip, C.K., Rezitis, J., Qi, Y., Bajaj, N., Koller, J., Farzi, A., Shi, Y.-C., Tasan, R., Zhang, L., Herzog, H.: Critical role of lateral habenula circuits in the control of stress-induced palatable food consumption. Neuron (2023)

[51] Comeras, L.B., Hörmer, N., Mohan Bethuraj, P., Tasan, R.O.: Npy released from gaba neurons of the dentate gyrus specially reduces contextual fear without affecting cued or trace fear. Frontiers in Synaptic Neuroscience 13, 635726 (2021)

[52] Schuman, B., Machold, R.P., Hashikawa, Y., Fuzik, J., Fishell, G.J., Rudy, B.: Four unique interneuron populations reside in neocortical layer 1. Journal of Neuroscience 39(1), 125–139 (2019)

[53] Chartrand, T., Dalley, R., Close, J., Goriounova, N.A., Lee, B.R., Mann, R., Miller, J.A., Molnar, G., Mukora, A., Alfiler, L., et al.: Morphoelectric and transcriptomic divergence of the layer 1 interneuron repertoire in human versus mouse neocortex. Science 382(6667), 0805 (2023)

[54] Boyle, M.P., Bernard, A., Thompson, C.L., Ng, L., Boe, A., Mortrud, M., Hawrylycz, M.J., Jones, A.R., Hevner, R.F., Lein, E.S.: Cell-type-specific consequences of reelin deficiency in the mouse neocortex, hippocampus, and amygdala. Journal of Comparative Neurology 519(11), 2061–2089 (2011)

[55] Shin, H., Adesnik, H.: Ndnf interneurons, spartans of the cortical column: Small in number, strong in impact. Neuron 109(13), 2041–2042 (2021)

[56] Meng, J.H., Schuman, B., Rudy, B., Wang, X.-J.: Mechanisms of dominant electrophysiological features of four subtypes of layer 1 interneurons. Journal of Neuroscience 43(18), 3202–3218 (2023)

[57] Kuang, X.-L., Zhao, X.-M., Xu, H.-F., Shi, Y.-Y., Deng, J.-B., Sun, G.-T.: Spatio-temporal expression of a novel neuron-derived neurotrophic factor (ndnf) in mouse brains during development. BMC neuroscience 11(1), 1–11 (2010)

[58] Lorenzetti, V., Costafreda, S.G., Rimmer, R.M., Rasenick, M.M., Marangell, L.B., Fu, C.H.: Brain-derived neurotrophic factor association with amygdala response in major depressive disorder. Journal of affective disorders 267, 103–106 (2020)

[59] Fernando, A.B., Murray, J.E., Milton, A.L.: The amygdala: securing pleasure and avoiding pain. Frontiers in behavioral neuroscience 7, 190 (2013)

[60] Wei, J.-a., Han, Q., Luo, Z., Liu, L., Cui, J., Tan, J., Chow, B.K., So, K.-F., Zhang, L.: Amygdala neural ensemble mediates mouse social investigation behaviors. National Science Review 10(1), 179 (2023)

[61] Oboti, L., Sokolowski, K.: Gradual wiring of olfactory input to amygdala feedback circuits. Scientific Reports 10(1), 5871 (2020)

[62] Manjila, S.B., Betty, R., Kim, Y.: Missing pieces in decoding the brain oxytocin puzzle: Functional insights from mouse brain wiring diagrams. Frontiers in Neuroscience 16, 1044736 (2022)

[63] Poller, W.C., Madai, V.I., Bernard, R., Laube, G., Veh, R.W.: A glutamatergic projection from the lateral hypothalamus targets vta-projecting neurons in the lateral habenula of the rat. Brain research 1507, 45–60 (2013)

[64] Penzo, M.A., Gao, C.: The paraventricular nucleus of the thalamus: an integrative node underlying homeostatic behavior. Trends in neurosciences 44(7), 538–549 (2021)

[65] Eichenbaum, H.: The hippocampus and mechanisms of declarative memory. Behavioural brain research 103(2), 123–133 (1999)

[66] Buzsáki, G., Moser, E.I.: Memory, navigation and theta rhythm in the hippocampal-entorhinal system. Nature neuroscience 16(2), 130–138 (2013)

[67] Çavdar, S., Özgur, M., Kuvvet, Y., Bay, H.H.: The cerebello-hypothalamic and hypothalamo-cerebellar pathways via superior and middle cerebellar peduncle in the rat. The Cerebellum 17, 517–524 (2018)

[68] Ulrich-Lai, Y.M., Herman, J.P.: Neural regulation of endocrine and autonomic stress responses. Nature reviews neuroscience 10(6), 397–409 (2009)

[69] Zeisel, A., Hochgerner, H., Lönnerberg, P., Johnsson, A., Memic, F., Van Der Zwan, J., Häring, M., Braun, E., Borm, L.E., La Manno, G., et al.: Molecular architecture of the mouse nervous system. Cell 174(4), 999–1014 (2018)

[70] Spatial_Visium. Mouse Brain Visium. https://support.10xgenomics.com/spatial-gene-expression/datasets/1.0.0/V1_Adult_Mouse_Brain. Accessed: 2023-11-06

[71] Castanza, A.S., Recla, J.M., Eby, D., Thorvaldsdóttir, H., Bult, C.J., Mesirov, J.P.: Extending support for mouse data in the molecular signatures database (msigdb). Nature Methods 20(11), 1619–1620 (2023)

[72] Chiareli, R.A., Carvalho, G.A., Marques, B.L., Mota, L.S., Oliveira-Lima, O.C., Gomes, R.M., Birbrair, A., Gomez, R.S., Simão, F., Klempin, F., et al.: The role of astrocytes in the neurorepair process. Frontiers in cell and developmental biology 9, 665795 (2021)

[73] Mills III, W.A., Woo, A.M., Jiang, S., Martin, J., Surendran, D., Bergstresser, M., Kimbrough, I.F., Eyo, U.B., Sofroniew, M.V., Sontheimer, H.: Astrocyte plasticity in mice ensures continued endfoot coverage of cerebral blood vessels following injury and declines with age. Nature Communications 13(1), 1794 (2022)

[74] Ross, J.M., Kim, C., Allen, D., Crouch, E.E., Narsinh, K., Cooke, D.L., Abla, A.A., Nowakowski, T.J., Winkler, E.A.: The expanding cell diversity of the brain vasculature. Frontiers in physiology 11, 600767 (2020)

[75] Pan, L., Trimarco, A., Zhang, A.J., Fujimori, K., Urade, Y., Sun, L.O., Taveggia, C., Zhang, Y.: Oligodendrocyte-lineage cell exocytosis and l-type prostaglandin d synthase promote oligodendrocyte development and myelination. Elife 12, 77441 (2023)

[76] Brockschnieder, D., Sabanay, H., Riethmacher, D., Peles, E.: Ermin, a myelinating oligodendrocyte-specific protein that regulates cell morphology. Journal of Neuroscience 26(3), 757–762 (2006)

[77] atlas.brain-map.org. http://atlas.brain-map.org (accessed: 27.01.2024)

[78] MacDonald, A., Lu, B., Caron, M., Caporicci-Dinucci, N., Hatrock, D., Petrecca, K., et al.: Single cell transcriptomics of ependymal cells across age, region and species reveals cilia-related and metal ion regulatory roles as major conserved ependymal cell functions. Front Cell Neurosci. 2021; 15: 703951 (2021)

[79] Cougnoux, A., Yerger, J.C., Fellmeth, M., Serra-Vinardell, J., Martin, K., Navid, F., Iben, J., Wassif, C.A., Cawley, N.X., Porter, F.D.: Single cell transcriptome analysis of niemann–pick disease, type c1 cerebella. International Journal of Molecular Sciences 21(15), 5368 (2020)

[80] Yuan, S., Norgard, R.J., Stanger, B.Z.: Cellular plasticity in cancer. Cancer discovery 9(7), 837–851 (2019)

[81] Griewank, K.G., Scolyer, R.A., Thompson, J.F., Flaherty, K.T., Schadendorf, D., Murali, R.: Genetic alterations and personalized medicine in melanoma: progress and future prospects. Journal of the National Cancer Institute 106(2), 435 (2014)

[82] Valdes-Mora, F., Handler, K., Law, A.M., Salomon, R., Oakes, S.R., Ormandy, C.J., Gallego-Ortega, D.: Single-cell transcriptomics in cancer immunobiology: the future of precision oncology. Frontiers in immunology 9, 2582 (2018)

[83] Yang, D., Jones, M.G., Naranjo, S., Rideout, W.M., Min, K.H.J., Ho, R., Wu, W., Replogle, J.M., Page, J.L., Quinn, J.J., et al.: Lineage tracing reveals the phylodynamics, plasticity, and paths of tumor evolution. Cell 185(11), 1905–1923 (2022)

[84] Bassiouni, R., Idowu, M.O., Gibbs, L.D., Robila, V., Grizzard, P.J., Webb, M.G., Song, J., Noriega, A., Craig, D.W., Carpten, J.D.: Spatial transcriptomic analysis of a diverse patient cohort reveals a conserved architecture in triple-negative breast cancer. Cancer Research 83(1), 34–48 (2023)

[85] Arora, R., Cao, C., Kumar, M., Sinha, S., Chanda, A., McNeil, R., Samuel, D., Arora, R.K., Matthews, T.W., Chandarana, S., et al.: Spatial transcriptomics reveals distinct and conserved tumor core and edge architectures that predict survival and targeted therapy response. Nature Communications 14(1), 5029 (2023)

[86] Bergholtz, H., Carter, J.M., Cesano, A., Cheang, M.C.U., Church, S.E., Divakar, P., Fuhrman, C.A., Goel, S., Gong, J., Guerriero, J.L., et al.: Best practices for spatial profiling for breast cancer research with the geomx® digital spatial profiler. Cancers 13(17), 4456 (2021)

[87] Wu, S.Z., Al-Eryani, G., Roden, D.L., Junankar, S., Harvey, K., Andersson, A., Thennavan, A., Wang, C., Torpy, J.R., Bartonicek, N., et al.: A single-cell and spatially resolved atlas of human breast cancers. Nature genetics 53(9), 1334–1347 (2021)

[88] Cancer Spatial. https://zenodo.org/record/4739739. Accessed: 2023-06-11

[89] Kim, H.K., Park, K.H., Kim, Y., Park, S.E., Lee, H.S., Lim, S.W., Cho, J.H., Kim, J.-Y., Lee, J.E., Ahn, J.S., et al.: Discordance of the pam50 intrinsic subtypes compared with immunohistochemistry-based surrogate in breast cancer patients: potential implication of genomic alterations of discordance. Cancer research and treatment: official journal of Korean Cancer Association 51(2), 737–747 (2019)

[90] Picornell, A., Echavarria, I., Alvarez, E., López-Tarruella, S., Jerez, Y., Hoadley, K., Parker, J., Monte-Millán, M., Ramos-Medina, R., Gayarre, J., et al.: Breast cancer pam50 signature: correlation and concordance between rna-seq and digital multiplexed gene expression technologies in a triple negative breast cancer series. BMC genomics 20(1), 1–11 (2019)

[91] Liu, L., Liu, L., Yao, H.H., Zhu, Z.Q., Ning, Z.L., Huang, Q.: Stromal myofibroblasts are associated with poor prognosis in solid cancers: a meta-analysis of published studies. PloS one 11(7), 0159947 (2016)

[92] Galvis-Jiménez, J.M., Curtidor, H., Patarroyo, M.A., Monterrey, P., Ramírez-Clavijo, S.R.: Mammaglobin peptide as a novel biomarker for breast cancer detection. Cancer biology & therapy 14(4), 327–332 (2013)

[93] Coleman, W.B.: Breast ductal carcinoma in situ: precursor to invasive brest cancer. The American journal of pathology 189(5), 942–945 (2019)

[94] Zhou, W., Jirström, K., Amini, R.-M., Fjällskog, M.-L., Sollie, T., Lindman, H., Sørlie, T., Blomqvist, C., Wärnberg, F.: Molecular subtypes in ductal carcinoma in situ of the breast and their relation to prognosis: a population-based cohort study. BMC cancer 13(1), 1–9 (2013)

[95] Allred, D.C., Wu, Y., Mao, S., Nagtegaal, I.D., Lee, S., Perou, C.M., Mohsin, S.K., O’Connell, P., Tsimelzon, A., Medina, D.: Ductal carcinoma in situ and the emergence of diversity during breast cancer evolution. Clinical cancer research 14(2), 370–378 (2008)

[96] Gini, C.: Memorie di Metodologia Statistica vol. 1. Libreria Goliardica, Padova, Italy (1955)

[97] Wright Muelas, M., Mughal, F., O’Hagan, S., Day, P.J., Kell, D.B.: The role and robustness of the gini coefficient as an unbiased tool for the selection of gini genes for normalising expression profiling data. Scientific reports 9(1), 17960 (2019)

[98] Kendall, M., Stuart, A., Ord, J.K.: Distribution Theory vol. 1. Arnold, London (1994)

[99] Sen, A.: Poverty, inequality and unemployment: Some conceptual issues in measurement. Economic and Political Weekly, 1457–1464 (1973)

[100] Giacomello, S., Salmén, F., Terebieniec, B.K., Vickovic, S., Navarro, J.F., Alexeyenko, A., Reimegård, J., McKee, L.S., Mannapperuma, C., Bulone, V., et al.: Spatially resolved transcriptome profiling in model plant species. Nature Plants 3(6), 1–11 (2017)

[101] Asp, M., Salmén, F., Ståhl, P.L., Vickovic, S., Felldin, U., Löfling, M., Fernandez Navarro, J., Maaskola, J., Eriksson, M.J., Persson, B., et al.: Spatial detection of fetal marker genes expressed at low level in adult human heart tissue. Scientific reports 7(1), 1–10 (2017)

[102] Moncada, R., Barkley, D., Wagner, F., Chiodin, M., Devlin, J.C., Baron, M., Hajdu, C.H., Simeone, D.M., Yanai, I.: Integrating microarray-based spatial transcriptomics and single-cell rna-seq reveals tissue architecture in pancreatic ductal adenocarcinomas. Nature biotechnology 38(3), 333–342 (2020)

[103] Maniatis, S., Äijö, T., Vickovic, S., Braine, C., Kang, K., Mollbrink, A., Fagegaltier, D., Andrusivová, Ž., Saarenpää, S., Saiz-Castro, G., et al.: Spatiotemporal dynamics of molecular pathology in amyotrophic lateral sclerosis. Science 364(6435), 89–93 (2019)

[104] McDavid, A., Finak, G., Yajima, M.: Mast: model-based analysis of single cell transcriptomics. Genome Biol 16(278), 10–1186 (2015)

[105] Hollander, M., Wolfe, D.A., Chicken, E.: Nonparametric Statistical Methods, 3rd edn. John Wiley & Sons, Hoboken, NJ (2013)

[106] Wilcoxon, F.: Individual comparisons by ranking methods. In: Breakthroughs in Statistics: Methodology and Distribution, pp. 196–202. Springer, New York, NY (1992)

[107] R Core Team: R: A Language and Environment for Statistical Computing. R Foundation for Statistical Computing, Vienna, Austria (2021). R Foundation for Statistical Computing. https://www.R-project.org/

